# Functional diversity of *NLRP3* gain-of-function mutants associated with CAPS autoinflammation

**DOI:** 10.1101/2023.09.22.558949

**Authors:** Camille Cosson, Romane Riou, Danish Patoli, Tingting Niu, Amaury Rey, Marine Groslambert, Elodie Chatre, Omran Allatif, Thomas Henry, Fabienne Venet, Florian Milhavet, Guilaine Boursier, Alexandre Belot, Yvan Jamilloux, Etienne Merlin, Agnès Duquesne, Gilles Grateau, Léa Savey, Alexandre Thibault Jacques Maria, Anne Pagnier, Solène Poutrel, Olivier Lambotte, Coralie Mallebranche, Frédéric Rieux-Laucat, Brigitte Bader-Meunier, Zahir Amoura, Isabelle Melki, Isabelle Touitou, Matthias Geyer, Sophie Georgin-Lavialle, Bénédicte F. Py

## Abstract

NLRP3-associated autoinflammatory disease (NLRP3-AID or CAPS) is an heterogenous group of monogenic autoinflammations associated with *NLRP3* gain-of-function mutations. The poor functional characterization of most *NLRP3* variants is a barrier to diagnosis although patients can be efficiently treated with anti-IL-1 approaches. In addition, while NLRP3 inflammasome is controlled by coordinated priming and activation signals, gain-of-functions of *NLRP3* variants have been only investigated in response to priming. Here, we functionally characterize 34 *NLRP3* variants *in vitro* by determining their activity in response to induction, priming and/or activation signals, and their sensitivity to inhibitors. We highlight the functional diversity of the gain-of-function mutants and describe four groups based on their profile of signals required for their activation, that correlate partly with the symptoms severities. We identify a new group of *NLRP3* mutants responding to the activation signal without priming, with patients often misdiagnosed. Our results identify key NLRP3 residues controlling the inflammasome activity and sensitivity to inhibitors. The comparison of four inhibitors on the 34 variants identifies inhibitory mechanisms with broader efficiency for future drug design. Altogether, our results provide new insights on NLRP3 activation and an explanatory mechanism for NLRP3-AID heterogeneity, and original tools for NLRP3-AID diagnosis and anti-inflammatory disease drug development.

**eTOC Summary:** Functional characterization of 34 CAPS-associated *NLRP3* variants identifies polymorphisms versus gain-of-function pathogenic mutants, and highlights diversities in the signals controlling their activation and in their sensitivity to inhibitors. This study provides tools for CAPS diagnosis and anti-inflammation drug development and insights on NLRP3 control mechanisms.

## Introduction

Cryopyrinopathies (Cryopyrin-associated periodic syndrome, CAPS), now referred as *NLRP3* mutation-associated autoinflammatory diseases (*NRLP3*-AID) were historically described as three dominant clinical entities: familial cold urticaria (FCAS), Muckle-Wells syndrome (MWS), and neonatal-onset multisystem inflammatory disease (NOMID, also referred to as chronic infantile neurological cutaneous and joint syndrome, CINCA).^1, 2^ All subtypes classically display cold-induced urticaria. FCAS was described as a benign form with mainly cutaneous features and arthralgia, whereas Muckle-Wells syndrome patients were described with urticaria, chronic inflammation and even recurrent fever, sensorineural deafness, ocular inflammation, headache, arthritis and could be complicated by inflammatory AA amyloidosis. NOMID begins at birth and is characterized by central nervous system inflammation such as chronic meningitis, skin involvement with a diffuse non-itchy urticarial rash, joint involvement including deforming arthropathy preferentially affecting the knees and facial dysmorphia characterized by the presence of frontal bumps and nasal saddle deformation. AA amyloidosis can complicate all forms of NLRP3-AID.

Most NLRP3-AID patients are highly responsive to IL-1-targeted therapies, especially on cutaneous, ocular and articular features and prevention of inflammatory amyloidosis.^2^ Nevertheless, the clinical response in patients with stable deafness, bone deformity, chronic renal failure is often poor, which is why early diagnosis is critical to prevent irrevocable damages. Beside clinical presentation, NLRP3-AID diagnosis mostly relies on genetics. However, classical genetic approaches, including analysis of familial segregation and recurrent association with the disease, are poorly efficient to distinguish the gain-of-function pathogenic *NLRP3* mutations from non-pathogenic variants. Indeed, NLRP3-AID is a rare disease which prevalence ranges between 1-3 per million. Despite some genotype-phenotype correlations, the symptoms and the severity may be variable between patients bearing identical mutations, even within one family. In addition, few low penetrance variants of *NLRP3* are associated with typical or atypical NLRP3-AID phenotypes, but also found in asymptomatic people. Finally, NLRP3-AID can be caused by somatic mosaicism, especially in NOMID often associated with *de novo* mutations.^3^ Of the 204 amino-acid substitutions or deletions in *NLRP3* described today, only 11% have been fully determined to be pathogenic (n=22) or benign (n=1), while 49% are likely pathogenic (n=96) or likely benign (n=4) and 40% are variants of uncertain significance (n=51, VUS), unsolved (n=6) or not classified (n=24).^4^

NLRP3 is a cytosolic stress sensor that upon activation assembles the inflammasome signaling complex controlling IL-1β and IL-18 secretion and pyroptosis, a pro-inflammatory cell death coupled with the release of many alarmins. NLRP3 is activated by a two-step mechanism engaged by coordinated priming and activation signals. Priming is usually provided by pattern recognition receptor (PRR) ligands, including the Toll-like receptor 4 (TLR4) agonist LPS, as well as cytokines. Priming was first described to increase the cellular level of key proteins in the pathway by transcriptional up-regulation of *NLRP3* itself, the inducible inflammasome substrate pro-IL-1β and many other inflammatory genes.^5^ Nevertheless, priming is now recognized to also trigger an ensemble of NLRP3 post-translational modifications that renders NLRP3 competent for activation.^6–8^ Activation signals correspond to diverse cell stresses, that target plasma membrane ion permeability (including the bacterial toxin nigericin), lysosomal rupture or mitochondrial functions for most of them. These signals converge towards a disruption of vesicular trafficking, which could be the common trigger for NLRP3 activation.^9^ Then, NLRP3 assembles an inflammasome complex controlling caspase-1 activation. Caspase-1 cleaves pro-IL-1β and pro-IL-18 into their mature forms, as well as Gasdermin D (GSDMD), which in turn forms pores in the plasma membrane, and ultimately mediates IL-1β/18 release and initiates pyroptosis.

Due to the aforementioned limitations of classical genetic approaches, the identification of gain-of-function *NLRP3* variants requires functional approaches. In addition, the diversity of symptoms and their genotype-correlation suggest that gain-of-function mutants may be heterogeneous in their activity, and that clinical manifestations may depend, at least in part, on the grade of gain-of-function. Current functional characterizations of *NLRP3* variants are largely based on the detection of IL-1β secretion in response to priming by LPS.^2^ Nevertheless, this approach does not distinguish between constitutively active and priming-activated mutants and does not detect mutations that would bypass the priming requirement.

We have developed a functional cell-based assay to screen for *NLRP3* variants that uncouples NLRP3 induction, priming and activation. For 34 *NLRP3* variants, we assessed pyroptosis and IL-1β/18 secretion from *NLRP3*-deficient U937 cells reconstituted with doxycycline-inducible NLRP3 variants in response to NLRP3 induction, priming and/or activation. The results were confirmed in primary monocytes from 15 patients carrying 9 different variants. These analyses efficiently discriminated gain-of-function mutants from polymorphisms without any impact on NLRP3 activity and evidenced the heterogeneity of the gain-of-function mutants that partly correlated with the severity of the symptoms. In particular, we evidenced a group of mutants responsive to the activation signal in the absence of prior priming that could not be detected in previous functional assays, leading to misdiagnosis of non-NLRP3-AID. Moreover, our study identified some key residues in the control of NLRP3 activity and sensitivity to inhibitors, as well as inhibitor mechanisms efficient towards the largest spectrum of *NLRP3* mutants.

## Results

### Reconstitution of *NLRP3*-deficient U937 human monocytes with inducible *NLRP3* variants associated with autoinflammation

In this study we analyzed 34 *NLRP3* variants identified in patients suffering auto-inflammation suggestive of NLRP3-AID (Table 1). 23 variants were selected among the most common variants associated with NLRP3-AID in France.^10^ 11 variants were included along the study upon the request of clinicians for diagnosis purpose. In order to compare the impact of *NLRP3* variants on inflammasome activity in a standardized genetic background, we used U937 human monocytes knocked-out for *NLRP3* by CRISPR/Cas9 and stably reconstituted them with doxycycline-inducible *NLRP3* variants (Figure 1, S1).^11^ The expression levels of *NLRP3* variants were similar or lower than the level of control WT *NLRP3* in reconstituted U937 cell lines treated with doxycycline, excluding that their putative gain-of-function may be caused by higher expression levels. In addition, NLRP3 levels in doxycycline-treated reconstituted U937 monocytes were similar to endogenous NLRP3 level in U937 cells treated with LPS, indicating that the system was physiologically relevant.

**Table 1:** *NLRP3* variants included in the study (Infevers database, personal observations and Louvrier et al.^3^). Frequencies based on exome sequencing of general population of all origins. *, patients with somatic mutation were asymptomatic. PYD, pyrin domain; NACHT, NAIP, CIITA, HET-E and TEP1 domain; LRR, Leucin-rich repeats domain; trLRR, transition LRR; FISNA, Fish-specific NACHT associated domain; NBD, nucleotide binding domain; WHD, winged helix domain; HD1/2, helical domain; VUS, variant of uncertain significance.

| Variants | Exon | Domain | Germinal/Somatic | Pathogenicity | Frequency |
| --- | --- | --- | --- | --- | --- |
| A75V | 2 | PYD | Germinal | VUS | 1e-04 |
| R168Q | 4 | NACHT (FISNA) | Germinal | Likely pathogenic | 1.991e-05 |
| V198M | 4 | NACHT (FISNA) | Germinal | VUS | 8.31e-03 |
| R260W | 4 | NACHT (NBD) | Germinal | Pathogenic | - |
| V262G | 4 | NACHT (NBD) | Germinal | Likely pathogenic | - |
| G301D | 4 | NACHT (NBD) | Germinal | Unsolved | - |
| D303H | 4 | NACHT (NBD) | Somatic | Likely pathogenic | - |
| D303N | 4 | NACHT (NBD) | Germinal | Pathogenic | - |
| E311K | 4 | NACHT (NBD) | Germinal | Likely pathogenic | - |
| I313V | 4 | NACHT (NBD) | Germinal | VUS | 2.1e-04 |
| I334V | 4 | NACHT (NBD) | Somatic | Likely pathogenic | - |
| T348M | 4 | NACHT (NBD) | Germinal/Somatic* | Pathogenic | - |
| A352V | 4 | NACHT (NBD) | Germinal | Pathogenic | - |
| L353P | 4 | NACHT (NBD) | Germinal | Likely pathogenic | - |
| K375E | 4 | NACHT (HD1) | Germinal | VUS | - |
| T436N | 4 | NACHT (WHD) | Germinal | Likely pathogenic | - |
| A439T | 4 | NACHT (WHD) | Germinal | Likely pathogenic | - |
| A439V | 4 | NACHT (WHD) | Germinal | Pathogenic | - |
| E525K | 4 | NACHT (HD2) | Germinal | Likely pathogenic | - |
| K565E | 4 | NACHT (HD2) | Germinal/Somatic* | VUS |  |
| E567G | 4 | NACHT (HD2) | Somatic | Likely pathogenic |  |
| K568N | 4 | NACHT (HD2) | Somatic | Likely pathogenic | - |
| G569R | 4 | NACHT (HD2) | Germinal | Pathogenic | - |
| Y570C | 4 | NACHT (HD2) | Germinal/Somatic | Likely pathogenic | - |
| E690K | 4 | trLRR | Germinal | Unsolved | - |
| M701T | 4 | trLRR | Germinal | VUS | 4.005e-6 |
| Q703K | 4 | trLRR | Germinal | VUS | 3.91e-2 |
| S726G/S896P | 5/8 | trLRR/LRR | Germinal | VUS | - |
| S726G | 5 | trLRR | Germinal | VUS | 5e-4 |
| G767S | 5 | LRR | Germinal | Likely pathogenic | - |
| Y859C | 7 | LRR | Germinal | Likely pathogenic | - |
| S896P | 8 | LRR | Germinal | VUS | 3.183e-5 |
| T952M | 9 | LRR | Germinal | VUS | 1.3e-3 |
| L1016F | 10 | LRR | Germinal | VUS | - |

**Figure 1.**
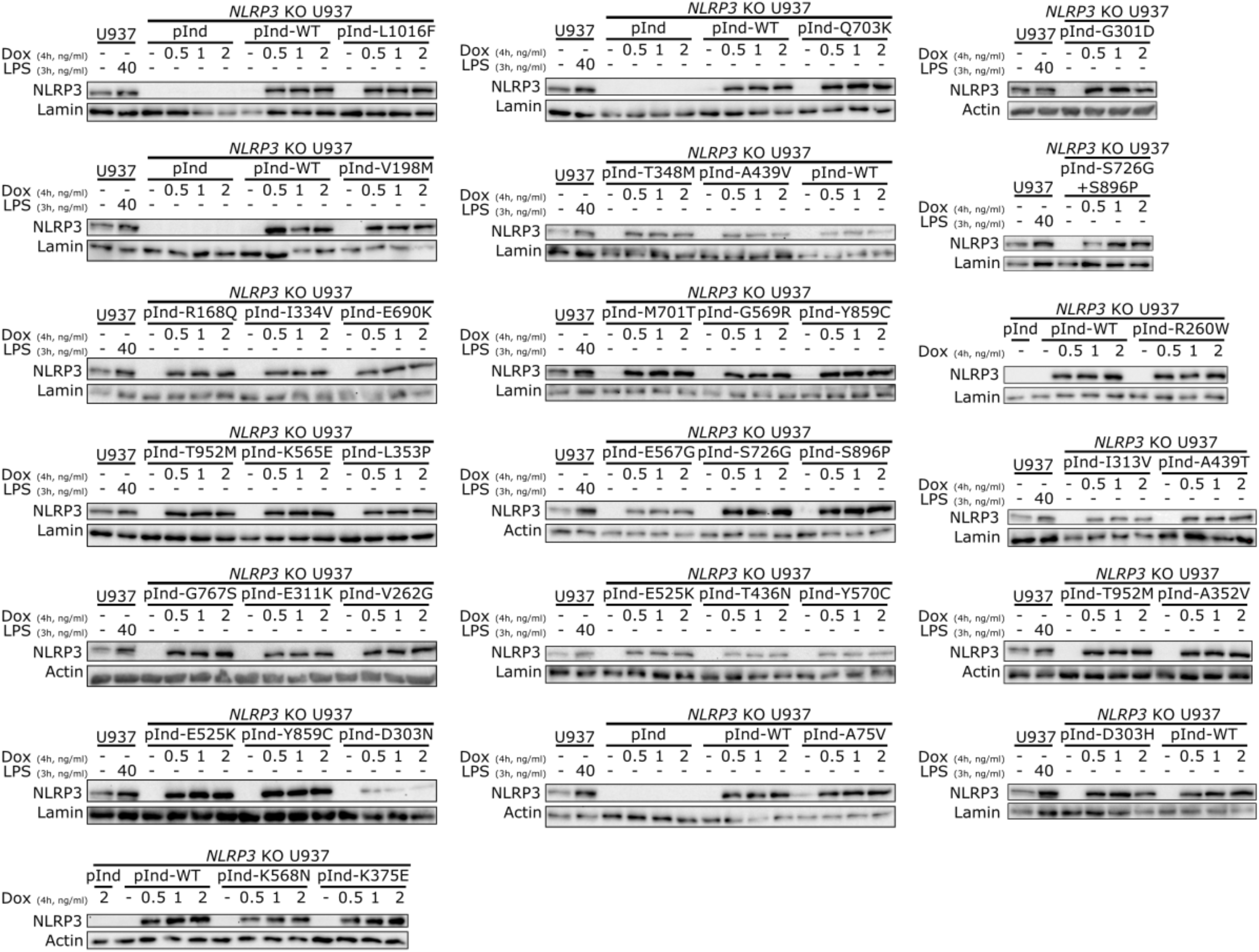
Reconstitution of *NLRP3*-deficient U937 with doxycycline-inducible *NLRP3* variants. *NLRP3*-deficient U937 reconstituted with doxycycline-inducible WT NLRP3 (pInd-WT), indicated *NLRP3* variants or the empty vector (pInd) were treated with doxycycline (indicated doses, 4h). U937 treated with LPS (40 ng/ml, 3h) were used as control. NLRP3 protein levels were analyzed by WB.

### Functional screen of *NLRP3* variants using pyroptosis as readout

The activities of NLRP3 variants were tested under (1) induction of their expression by doxycycline (Dox), (2) expression induction followed by LPS as a priming signal (Dox+LPS), (3) expression induction followed by nigericin as an activation signal (Dox+Nig) and (4) expression induction followed by both priming and activation signals (Dox+LPS+Nig) (Figure S1). We first assessed NLRP3 activity by using pyroptosis as a readout (Figure 2A). We measured propidium iodide incorporation over time in Hoechst-counterstained cells using high content screening (HCS) microscopy (Figures 2, S2). Cell death percentages over time for each variant and following each treatment were compared to those of U937 cells reconstituted with WT *NLRP3* by fitting a linear mixed model (Figure 3). *NLRP3* variants could be classified in 5 functional groups based on their activation upon induction, priming and/or activation signals. U937 cells expressing group#1 *NLRP3* variants, as for example K568N, underwent cell death upon doxycycline-mediated induction of *NLRP3* expression (Figures 2B, 3). Group#1 variants corresponded therefore to constitutively active mutants. U937 cells expressing group#2 *NLRP3* variants, as E311K, underwent cell death upon NLRP3 expression and, either priming with LPS or activation with nigericin (Figures 2C, 3). U937 cells expressing group#3 *NLRP3* variants, as A439V, underwent cell death upon NLRP3 expression and priming with LPS (Figures 2D, 3). U937 cells expressing group#4 *NLRP3* variants, as R168Q, underwent cell death upon NLRP3 expression and activation with nigericin (Figures 2E, 3). Finally, U937 cells expressing group#5 *NLRP3* variants, as T952M, underwent cell death upon NLRP3 expression, priming with LPS and activation with nigericin (Figures 2F, 3). Group#5 variants corresponded to variants with no gain-of-function in reconstituted U937 cells. In our experimental scheme, cell death monitoring was initiated when nigericin was added, *i.e.* 3h after doxycycline treatment. In order to investigate early cell death for group#1 *NLRP3* variants, we imaged propidium iodide incorporation immediately following doxycycline treatment (Figure S3). Permeabilization of U937 expressing group#1 *NLRP3* variants was detectable starting from 2.5h following doxycycline treatment, validating our experimental scheme.

**Figure 2.**
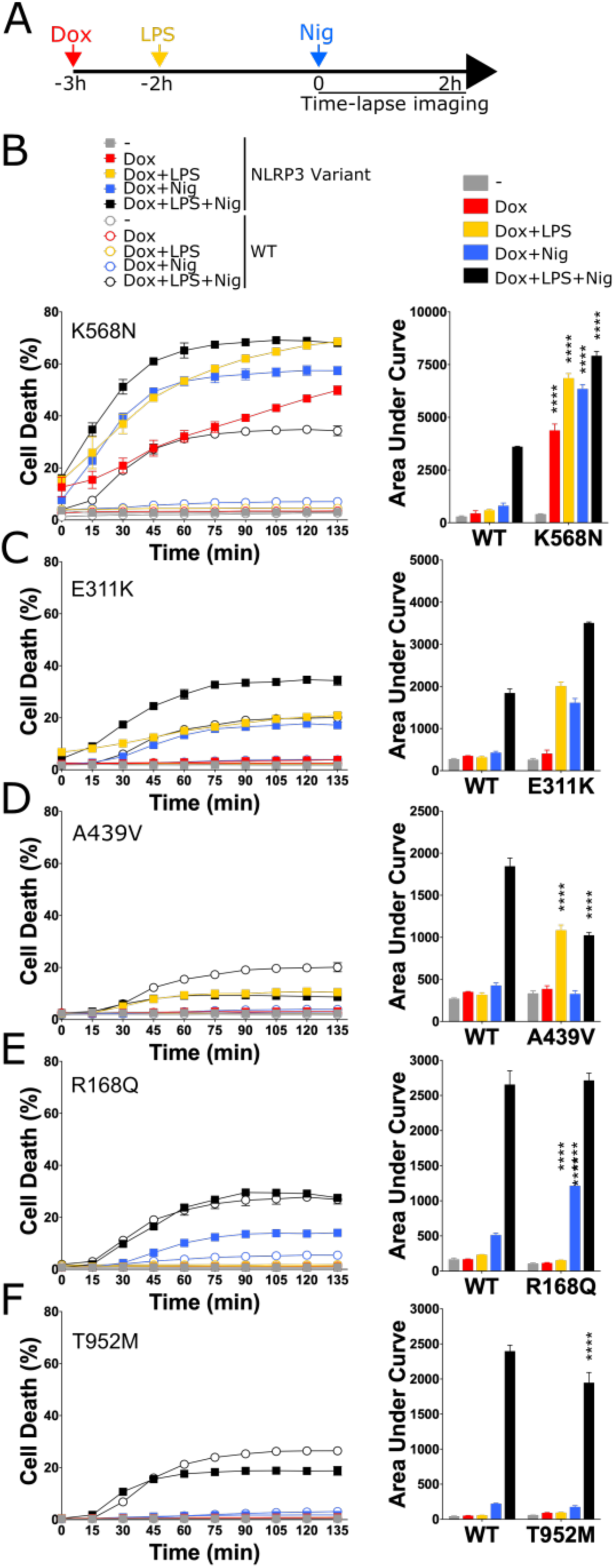
NLRP3 expression, priming and/or activation triggers pyroptosis in reconstituted U937 depending on the *NLRP3* variants. *NLRP3*-deficient U937 cells reconstituted with doxycycline-inducible *NLRP3* variants were treated with doxycycline (1 μg/ml, 3h), LPS (40 ng/ml, 2h) and nigericin (15 μg/ml) before cell death was monitored by PI incorporation over time quantified by time-lapse high content microscopy (A-F). B, K568N (group#1). C, E311K (group#2). D, A439V (group#3). E, R168Q (group#4). E, T952M (group#5). Means of duplicates and 1 SD (left panel) and means of area under the curve of duplicates and 1SD (right panel) are represented. Two-way ANOVA multiple comparisons of each variant with WT control with corresponding treatment, ****, p <0.0001. One experiment done in duplicates representative of 2-8 independent experiments is shown. Results obtained with one variant typical of each functional group are represented. Results obtained with all tested variants are presented in Figure S2. Statistical analysis including all independent experiments are represented in Figure 3.

**Figure 3.**
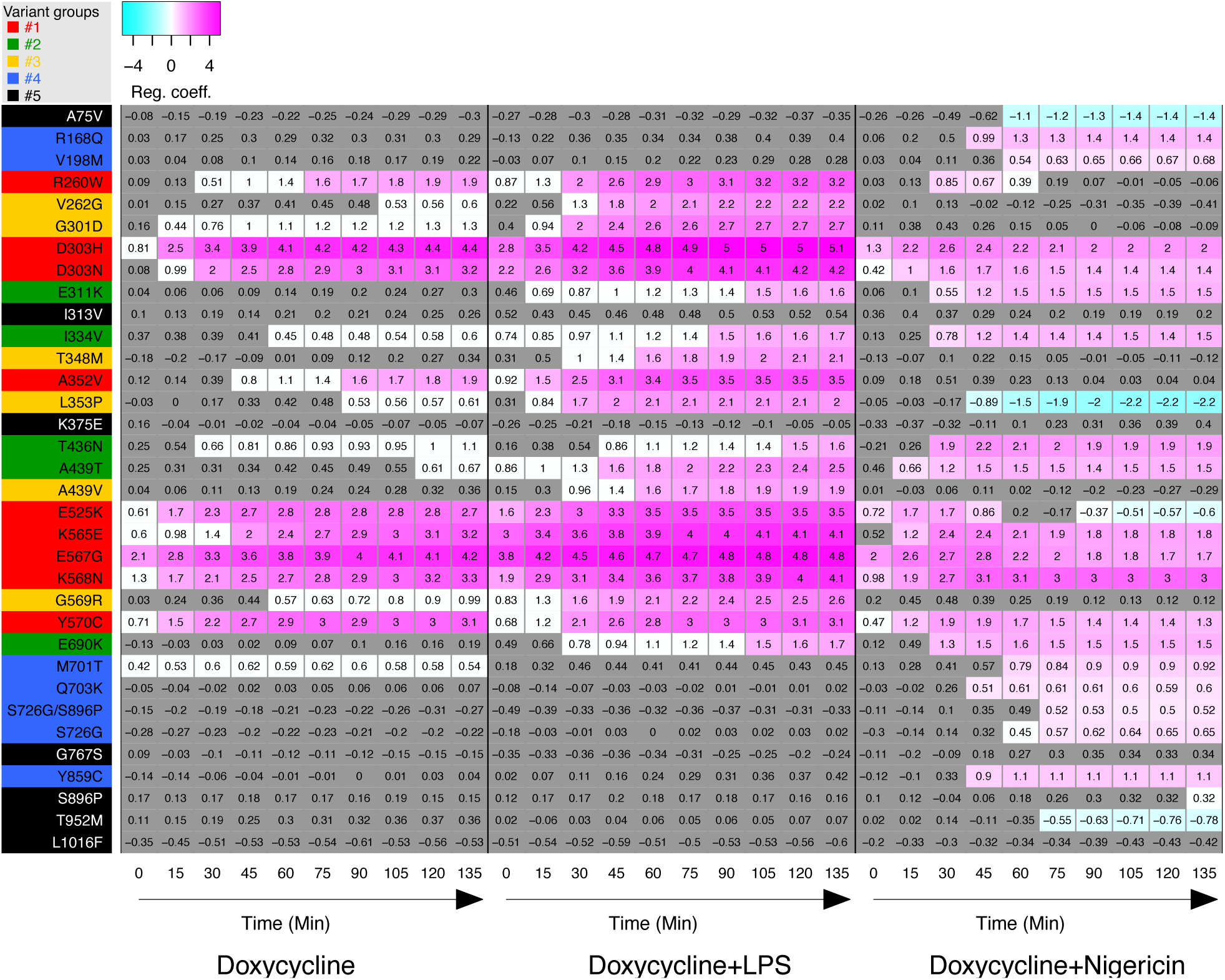
Relative cell death of U937 expressing *NLRP3* variants as compared to U937 expressing WT *NLRP3* in response to NLRP3 expression, priming and/or activation. Regression coefficient (RC) heatmap from the generalized linear mixed model for each variant as compared to the WT. Positive RC denotes increased cell death in the considered variant as compared to the WT, and conversely. RC corresponding to statistically significant increases (magenta) or decreases (cyan) in cell death are color-coded (p<0.05). Not significant, p>0.05 (grey). Based on the results, variants are classified in 5 functional groups: constitutive active variants (group#1, red), variants active upon either priming or activation signal (group#2, green), variants active upon priming signal (group#3, yellow), variants active upon activation signal (group#4, blue), and mutants active upon priming and activation signals (no gain-of-function, group#5, black).

### Functional screen of *NLRP3* variants using IL-18 and IL-1β secretions as readouts

As a complementary approach, we assessed NLRP3 activity by measuring secretion of inflammasome-dependent cytokines, *i.e.* IL-18 and IL-1β, following NLRP3 induction, priming and activation. LPS priming did not induce pro-IL-1β expression in U937 monocytes, while pro-IL-1β was induced in PMA-differentiated U937 (Figure 4A). As expected, pro-IL-18 was expressed in basal conditions in U937, and not increased upon PMA differentiation and LPS priming. We therefore measured IL-18 and IL-1β secretion in U937 macrophages differentiated with PMA 50 ng/ml for 16h, followed by treatments with doxycycline, LPS and/or nigericin (Figures 4B-H). TNF was measured as a control for NLRP3-independent cytokine secretion. IL-18, IL-1β and TNF secretions for each variant following each treatment were compared to those of U937 cells reconstituted with WT *NLRP3* by realizing linear mixed-effects models (Figures 5, S5). Notably, PMA differentiation was associated with low expression of pro-IL-1β in reconstituted U937 *NLRP3* KO cells in the absence of LPS priming and lower induction of *NLRP3* following doxycycline treatment as compared with undifferentiated cells (Figure 4C).

**Figure 4.**
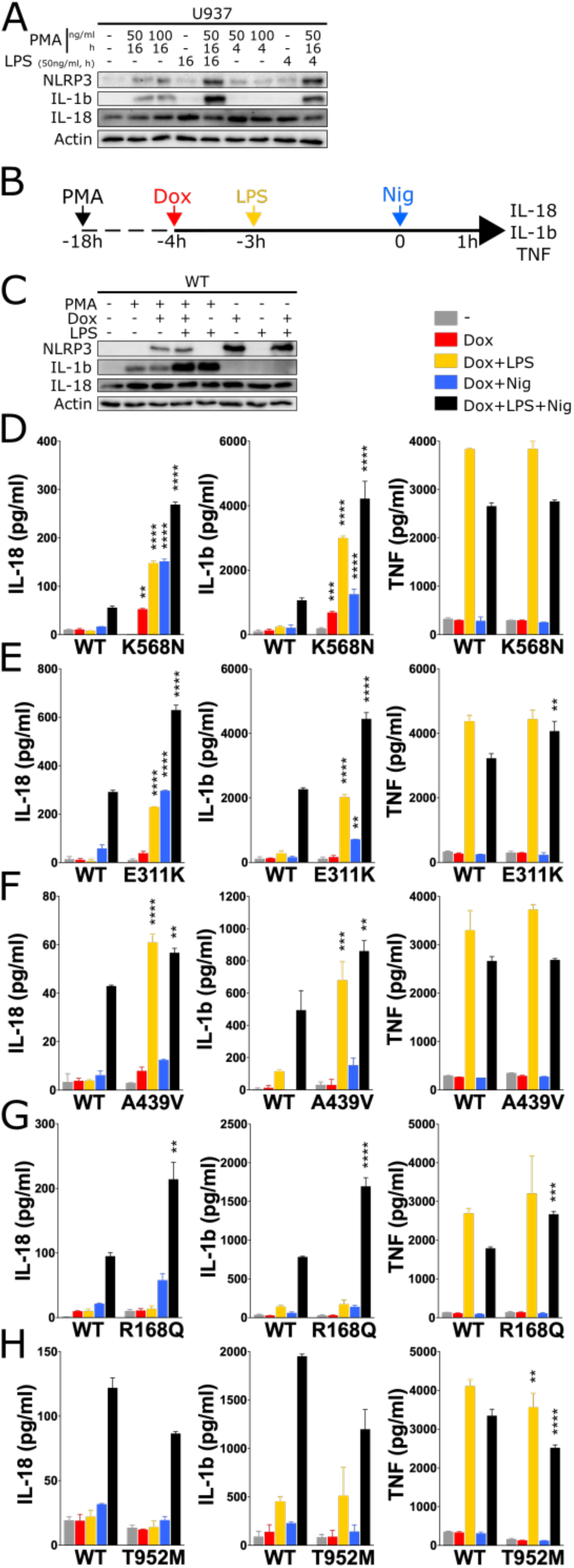
NLRP3 expression, priming and/or activation triggers IL-18 and IL-1β secretion in reconstituted U937 depending on the *NLRP3* variants. A, U937 cells were differentiated in PMA (50 or 100ng/ml, 4 or16h) and treated with LPS (50 ng/ml, 4-16h). Protein levels of pro-IL-18, pro-IL-1β and NLRP3 were assessed by WB. Actin is used as loading control. B-H, *NLRP3*-deficient U937 cells reconstituted with doxycycline-inducible *NLRP3* variants were differentiated in PMA (50 ng/ml) and treated with doxycycline (1 μg/ml), LPS (40 ng/ml) and nigericin (15 μg/ml) as indicated. NLRP3, pro-IL-1β, pro-IL-18 and actin levels (as control) were assessed by WB in lysates of *NLRP3*-deficient U937 cells reconstituted with doxycycline-inducible WT *NLRP3* (C). IL-18, IL-1β and TNF levels (as control) were assessed in cell supernatant by ELISA in K568N (group#1) (D), E311K (group#2) (E), A439V (group#3) (F), R168Q (group#4) (G) and T952M (group#5) (H). Means of duplicates and 1 SD are represented. Two-way ANOVA multiple comparisons of each variant with WT control with corresponding treatment on the same plate, *, p <0.05; **, p <0.01; ***, p <0.001; ****, p <0.0001. One experiment done in duplicates representative of 3-18 independent experiments is shown. Results obtained with one variant typical of each functional group are represented. Results obtained with all tested variants are presented in Figure S4. Statistical analysis including all independent experiments are represented in Figure 5.

**Figure 5.**
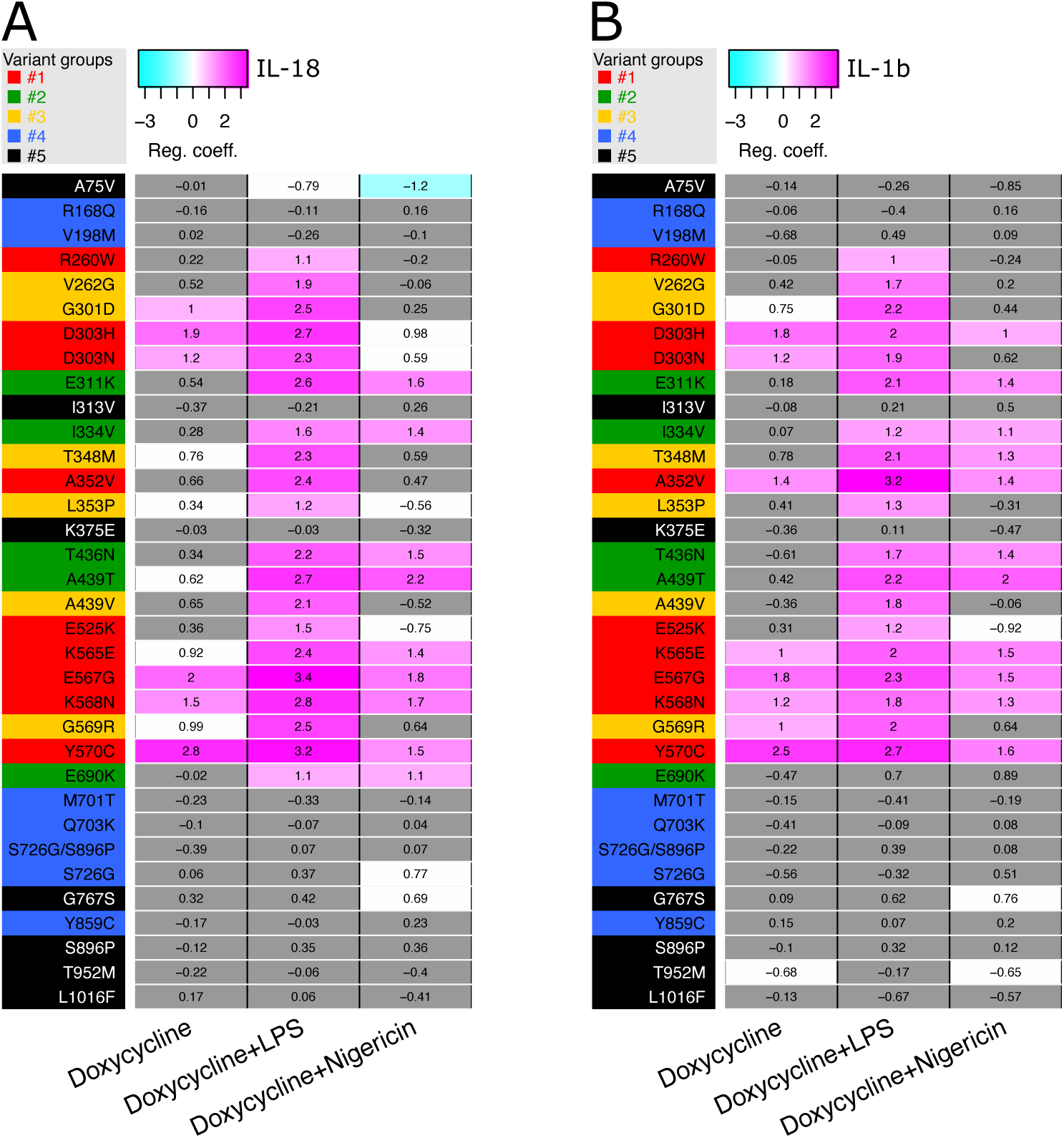
Relative IL-18 and IL-1β secretion of U937 cells expressing *NLRP3* variants as compared to U937 cells expressing WT *NLRP3* in response to NLRP3 expression, priming and/or activation. Regression coefficient (RC) heatmap from the lmm modeling for each variant as compared to the WT. Positive RC denotes increased IL-18 (A) and IL-1β (B) concentrations by the variant as compared to *NLRP3* WT, and conversely. RC corresponding to statistically significant increases (magenta) or decreases (cyan) in cytokine secretions are color-coded (p<0.05 and, RC>+1 or RC<-1 respectively). Not significant, p>0.05 (grey).

For all variants IL-1β secretion was higher in magnitude than IL-18 secretion, and constituted a more reproducible readout. For most *NLRP3* variants, IL-18 and IL-1β secretion matched induction of pyroptosis in response to NLRP3 expression, priming and activation signals by reconstituted U937. Cells expressing group#1 *NLRP3* variants, as for example K568N, secreted IL-18 and IL-1β in response to doxycycline (Figures 4D, 5). U937 cells expressing group#2 *NLRP3* variants, as for example E311K, secreted IL-18 and IL-1β in response to either LPS priming or activation with nigericin (Figures 4E, 5). U937 cells expressing group#3 *NLRP3* variants, as for example A439V, secreted IL-18 and IL-1β in response to LPS priming (Figures 4F, 5). U937 cells expressing group#4 *NLRP3* variants, as for example R168Q, secreted IL-18 in response to nigericin activation signal (Figures 4G, 5). Finally, U937 cells expressing group#5 *NLRP3* variants, as for example T952M, secreted IL-18 and IL-1β only in response to priming and activation signals (Figures 4H, 5).

In LPS-primed conditions, the statistical test robustly detected increased IL-18/1β secretion for all variants of the group#1, group#2 and group#3 as compared to U937 cells expressing WT *NLRP3* (Figure 5). Nevertheless, in the absence of LPS priming, several discrepancies could be noted between IL-18/1β secretion and pyroptosis. U937 cells expressing group#1 *NLRP3* variants R260W and E525K did not robustly secrete IL-18/1β in response to doxycycline, while they underwent pyroptosis. Similarly, U937 cells expressing group#1 *NLRP3* variants R260W, D303N and E525K as well as U937 cells expressing all group#4 *NLRP3* variants did not robustly secrete IL-18/1β in response to doxycycline and nigericin, while they underwent pyroptosis. We might speculate that lower NLRP3 expression in PMA-differentiated U937 in response to doxycycline and low expression of pro-IL-1β in these unprimed conditions may explain the lower sensitivity of the IL-18/1β secretion readout. On the opposite, U937 expressing group#3 *NLRP3* variant T348M secreted IL-1β in response to doxycycline and nigericin, indicating that this variant may exhibit partial activity that may lead to IL-18 and IL-1β secretion without cell death in this condition. As a control, U937 cells reconstituted with *NLRP3* variants secreted similar amount of TNF as U937 cells reconstituted with WT *NLRP3* in all conditions (Figure S5).

### Sensitivity of *NLRP3* variants to NLRP3 chemical inhibitors

We next investigated the sensitivity of the *NLRP3* variants to inflammasome inhibitors. We tested direct NLRP3 inhibitors MCC950 and CY-09.^12, 13^ MCC950 interacts with NLRP3 into a cleft in between the NACHT and the LRR domains in proximity to the ATP binding site and locks NLRP3 in an inactive conformation.^14–17^ CY-09 binding to NLRP3 depends on the integrity of NLRP3 ATP binding site and its inhibitory mechanism remains debated.^13, 15^ We also tested two indirect inhibitors: the deubiquitinase inhibitor G5, which impairs NLRP3 deubiquitination by BRCC3,^18^ and the protein kinase D inhibitor CRT0066101, which impairs NLRP3 phosphorylation at the Golgi during the activation process.^19^ We determined the optimal dose of each inhibitor using *NLRP3*-deficient U937 reconstituted with WT *NLRP3* treated with doxycycline, LPS and nigericin to reach similar 80% inhibition for all tested inhibitors (Figure S6). We then measured cell death following Dox (group#1), Dox+LPS (groups#1-3), Dox+Nig (groups#1, 2, 4), or Dox+LPS+Nig (groups#1-5) in the presence of these inhibitors added before the last treatment for each condition (Figures 6, S7-11). For each variant and for each treatment, we compared cell death in the presence of each of the inhibitors to vehicle only by fitting a linear mixed-effects model to AUC value databases (Figure 7). None of the group#5 variants showed reduced sensitivity to any of the inhibitors, confirming the absence of gain-of-function of these variants.

**Figure 6.**
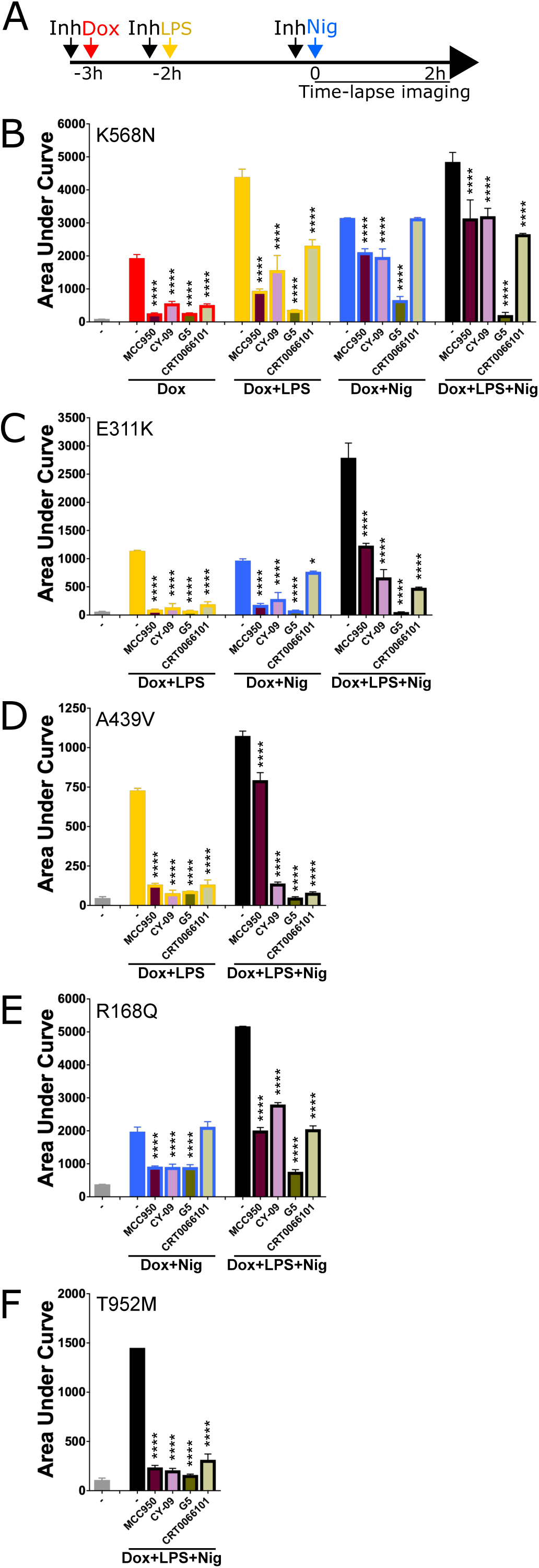
Cell death of U937 expressing *NLRP3* variants in response to NLRP3 expression, priming and/or activation in presence of NLRP3 inhibitors. A-F. *NLRP3*-deficient U937 cells reconstituted with doxycycline-inducible *NLRP3* variants were treated with doxycycline (1 μg/ml, 3h), LPS (40 ng/ml, 2h) and nigericin (15 μg/ml) in the presence of MCC950 (1 μM), CY-09 (50 μM), G5 (1 μM) and CRT006101 (0.5 μM) NLRP3 inhibitors, or DMSO vehicle, (added 20 min before the last treatment) and cell death was monitored by PI incorporation over time quantified by time-lapse high content microscopy for 2h (A). B, K568N (group#1). C, E311K (group#2). D, A439V (group#3). E, R168Q (group#4). F, T952M (group#5). Means of area under the curve of duplicates and 1SD are represented. Two-way ANOVA multiple comparisons of each inhibitor with the corresponding treatment with vehicle only. *, p <0.05; **, p <0.01; ***, p <0.001; ****, p <0.0001. One experiment done in duplicates representative of 2 independent experiments is shown. Results obtained with one variant typical of each functional group are represented. Results obtained with all tested variants are presented in Figure S7-11. Statistical analysis including all independent experiments are represented in Figure 7.

**Figure 7.**
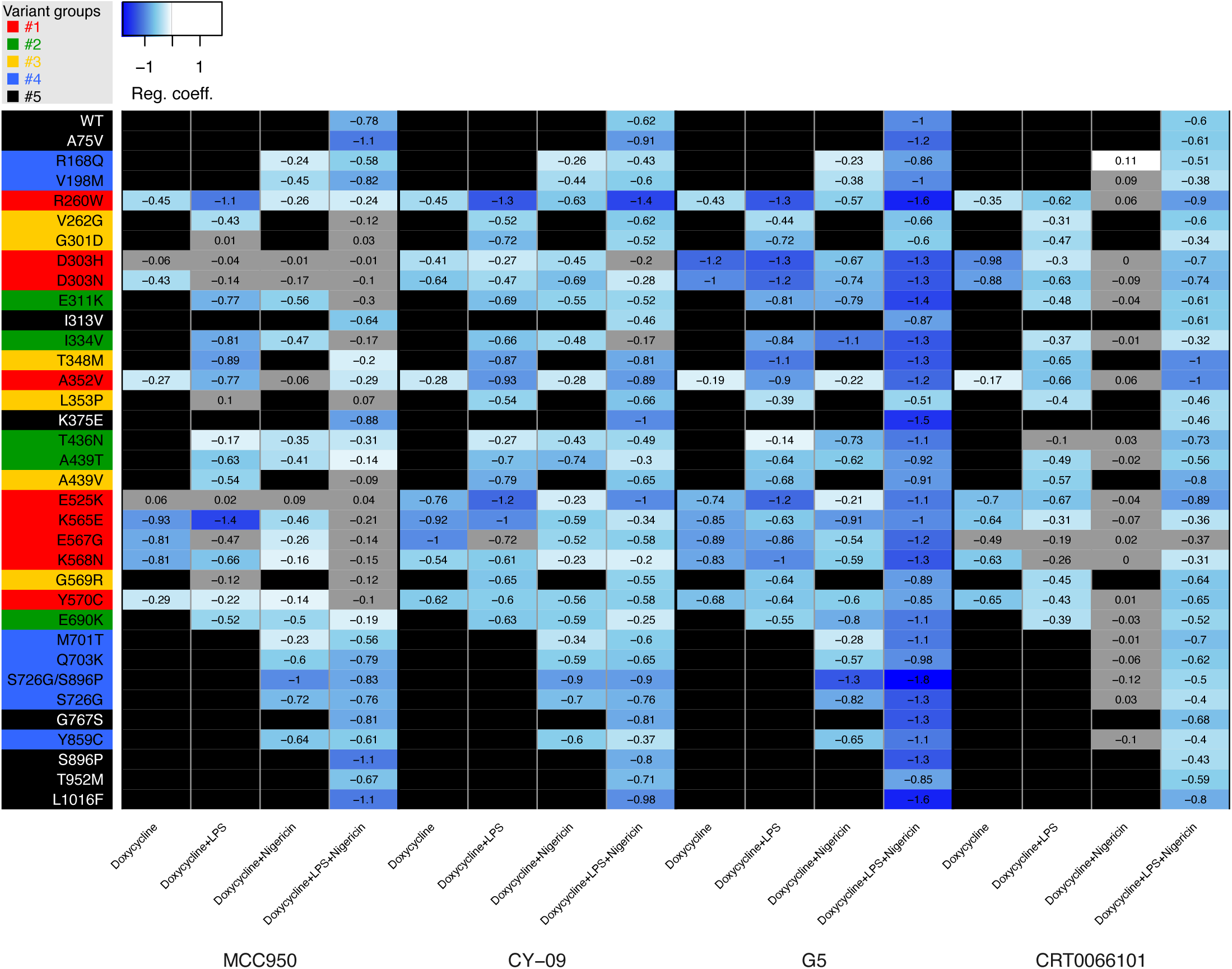
Inhibition of cell death of U937 expressing each *NLRP3* variants in response to NLRP3 expression, priming and/or activation in presence of NLRP3 inhibitors. Regression coefficient (RC) heatmap from the linear mixed model of PI incorporation AUC for each *NLRP3* variant with a specific treatment for each inhibitor, as compared to vehicle only. Negative coefficient denoted inhibition of cell death, and conversely. For significant values, RC <-0.1 are color-coded in blue. Not significant, p>0.05 (grey). Variant*treatment combinations that do not trigger cell death were not tested and are represented in black.

Most gain-of-function mutants are sensitive to MCC950, including all the mutants of group#4. Nevertheless, consistent with previous studies, several *NLRP3* mutants were resistant.^12, 17, 20^ D303H and E525K (group#1) as well as G301D, L353P and G569R (group#3) were resistant to MCC950 independently of the signals (priming or activation) provided. These mutations may affect NLRP3 ability to bind to MCC950, or impair the ability of MCC950 to lock NLRP3 in a closed conformation (Figure S13A, D). I334V, A439T and E690K (group#2), and V262G, A439V and T348M (group#3) were sensitive to MCC950 when added before the priming signal, but were resistant to MCC950, when added on primed U937 before the activation signal. In this last condition, these mutants may switch to an active conformation upon priming before the addition of MCC950. Similarly, K565E, E567G, K568N and Y570C (group#1) were resistant to MCC950, when added before the activation signal on primed cells. Noteworthy, K568N and Y570C were also partly resistant to MCC950, when added before the activation signal on non-primed cells.

E311K, I334V, A439T and E690K (group#2) were inhibited by MCC950 when added before priming or activation signal on non-primed cells, while T436N (group#2) was resistant to MCC950 when added before the priming signal, but sensitive to MCC950 when added before the activation signal. This suggested that while both priming and activation signals resulted in active T436N form, the structural switches underlying this activation process may differ. R260W (group#1) was inhibited by MCC950 when added before the LPS priming signal, or before the activation signal on primed cells. Although MCC950 was much less potent in this last condition, this may indicate that the LPS-induced structural switch in these mutants was reversible and that MCC950 may lock back NLRP3 in a closed conformation. D303N (group#1) was sensitive to MCC950 when added before its protein induction, but resistant if added later before the priming or the activation signal, suggesting that upon induction D303N may transiently adopt a closed conformation stabilized by MCC950, but quickly switch to an irreversible active conformation which may be stabilized by its oligomerization. In contrast to MCC950, no *NLRP3* variants showed strong resistance to the direct inhibitor CY-09. E525K, K568N (group#1) and R168Q (group#4) showed partial resistance to CY-09 when added before the activation signal to non-primed cells. D303H (group#1) and T436N (group#2) showed partial resistance when CY-09 was added before the priming signal. A352V (group#1) showed partial resistance when CY-09 was added upon induction or before the activation signal.

Concerning the indirect inhibitors, no *NLRP3* variants showed strong resistance to the deubiquitinase inhibitor G5. E525K (group#1) and R168Q (group#4) showed partial resistance when G5 was added before the activation signal to non-primed cells. A352V (group#1) was partially resistant when G5 was added before the induction or the activation signal. T436N (group#2) was partially resistant when G5 was added before the priming signal. CRT0066101 was inefficient in inhibiting any mutant of groups#1, 2 or 4 when added before the activation signal in non-primed cells. In addition, E567G (group#1) and T436N (group#2) were resistant to CRT0066101 when added before priming, and A352V (group#1) was partially resistant to CRT0066101 when added upon induction. Noteworthy, observed resistance to all inhibitors of A352V (if added before induction or activation of unprimed cells), T436N (if added before priming) and E525K (if added before activation of unprimed cells) may be artifactual as these mutants showed the weakest gain-of-function of their groups in these conditions (Figure 3).

### Activity of *NLRP3* variants in patient primary monocytes

In addition, we developed a functional assay on primary monocytes adapted for small volume pediatric blood samples as it requires less than 0.2x10^6^ monocytes. We assessed pyroptosis of primary monocytes from 15 patients bearing *NLRP3* variants in response to LPS and/or nigericin, and its sensitivity to MCC950 inhibition (Figures 8A). Priming and activation signals have both been independently described to be dispensable for NLRP3 activation in human monocytes.^21, 22^ In our experimental settings (dose, kinetics), LPS priming alone did not cause pyroptosis in monocytes from healthy donors (Figure S12). In the absence of priming, nigericin triggered pyroptosis in a MCC950-sensitive manner although with slower kinetics than in the presence of priming (Figure S12). In this context, additional priming signals from death of neighboring cells upon transportation or monocytes purification procedures could not be excluded.

**Figure 8.**
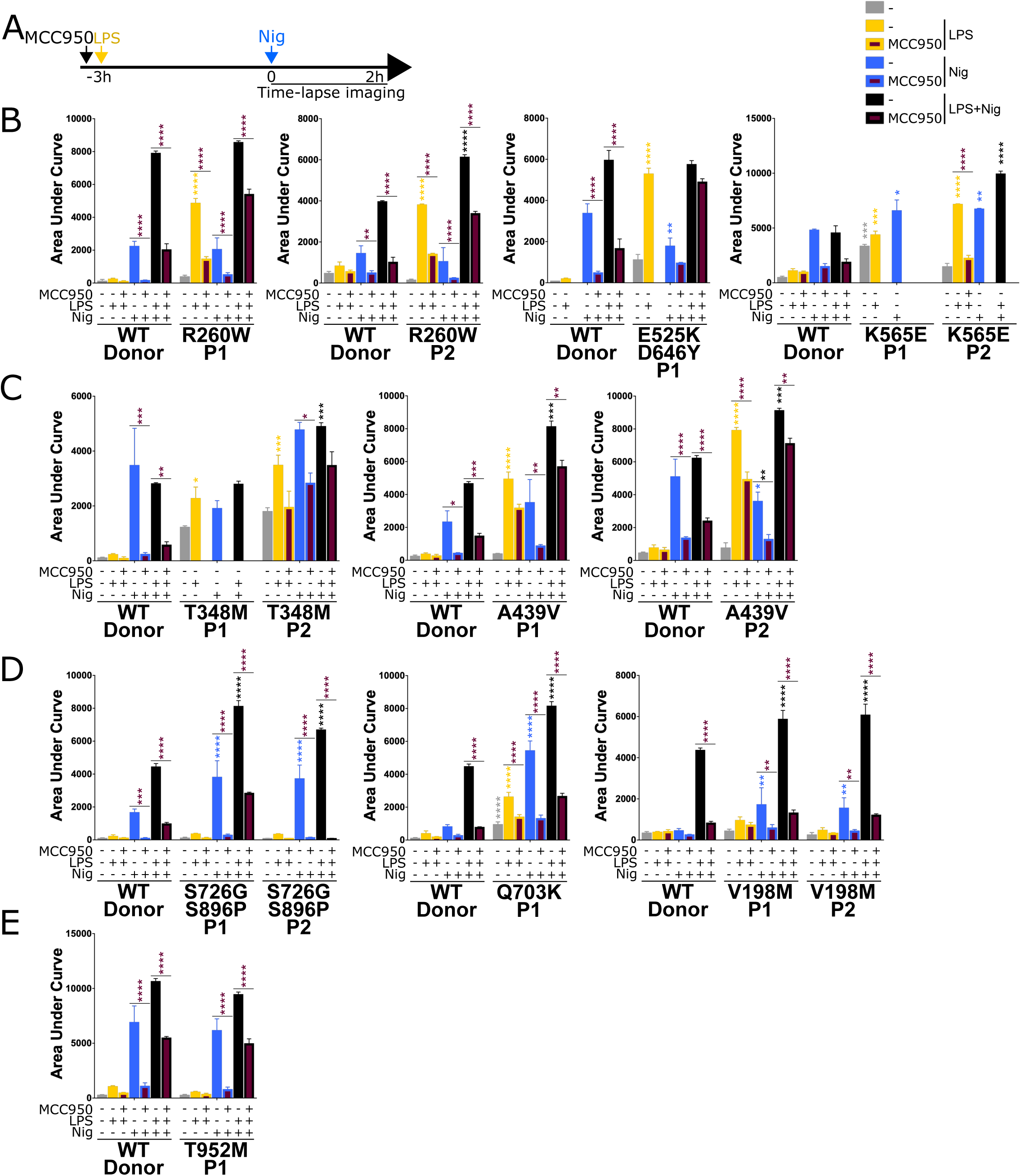
Patient monocytes respond differently to NLRP3 priming and/or activation, and NLRP3 inhibitor MCC950 depending on *NLRP3* variants. A, Monocytes of 15 autoinflammatory patients and healthy donors (WT) were treated with LPS (40 ng/ml, 3h) and nigericin (5 μg/ml) in the presence of MCC950 (1 μM, added 15 min before LPS) and cell death was monitored by PI incorporation over time quantified by time-lapse high content microscopy for 2h. B, R260W, E525K+D646Y and K565E (group#1). C, T348M and A439V (group#3). D, S726G+S896P, Q703K and V198M (group#4). E, T952M (group#5). Means of area under the curve of 2-6 replicates and 1SD are represented. Two-way ANOVA multiple comparisons of each variant vs WT control with corresponding treatment, and in the presence of MCC950 vs control with corresponding genotype*treatment in the absence of MCC950, ****, p <0.0001, ***, p <0.001, **, p <0.01, *, p <0.05. One experiment done in 2-6 replicates is shown. Data represented in the same graph correspond to one same experiment on relatives. Missing data correspond to conditions that were not tested due to insufficient monocyte numbers.

NLRP3 induction and priming could not be discriminated in this experimental setting, and monocytes from patients bearing *NLRP3* variants of group#1 (R260W, E525K+D646Y and K565E) and group#3 (T348M and A439V) showed increased pyroptosis in response to LPS as compared to their respective control monocytes from healthy donors (Figures 8B, C). Monocytes from patients bearing *NLRP3* variants of group#4 (S726G+S896P, Q703K and V198M) showed increased pyroptosis in response to nigericin (Figure 8D). Monocytes from a patient bearing a *NLRP3* variant of group#5 (T952M) showed no gain-of-function relative to their controls (Figure 8E). MCC950 inhibited pyroptosis of all but monocytes from patient bearing E525K (Figures 8 B-E), consistent with inhibition assays on reconstituted U937 cells (Figure 7).

## Discussion

NLRP3-AID clinical manifestations are diverse and most of them are poorly specific. Diagnosis relies largely on the identification of *NLRP3* variants by genetic analysis, but only 11% of known variants have been fully characterized as benign or pathogenic (Infevers 29/05/2023), while the functional link to the disease remains to be established for all the others.^4^ In addition, gain-of-functions of *NLRP3* variants have been assessed by monitoring NLRP3 activity in response to priming signal, which does not discriminate constitutive from priming-induced activity, and NLRP3 activity in response to activation signals in the absence of priming has not been considered.^23^ By reconstitution of *NLRP3*-deficient U937 monocytes, we functionally characterized 34 *NLRP3* variants, including the most frequent in France, on an identical genetic background. For 9 variants, results were confirmed on primary monocytes from 15 patients. We evidenced the functional diversity of *NLRP3* variants which we classified in 5 groups. Identification of *NLRP3* variants with no gain-of-function (group#5) will exclude NLRP3-AID diagnosis and guide decision toward broader genetic analysis for patients positive for these variants after gene panel approaches. Genotype-phenotype correlation are difficult to assess. Indeed, severities vary between patients bearing identical mutations, and in the anti-IL-1 therapy era, delayed access to treatment rather than the genotype determines symptoms severity. Nevertheless, the low genotype overlap between somatic and germline mutations suggest that somatic mutations, that may be responsible for 0.5%-19% of NLRP3-AID cases, would be highly detrimental and incompatible with life if germline. Interestingly, the functional diversity identified in our study strongly correlates with somatic vs germline mutation. All somatic mutations included in our study (n=5) correspond to constitutively active mutants (group#1, n=4) or mutants activated by priming or activation signal (group#2, n=1). 44% of group#1 mutants are associated with mosaicism (4/9), 20% of group#2 (1/5), but none of groups#3-4 (0/6, each).

Comparison of the sensitivity of *NLRP3* mutants to two direct and two indirect inhibitors evidenced that direct inhibitor CY-09 was efficient on more variants than MCC950, and that the deubiquitinase inhibitor G5 was efficient on all mutants. This suggests that refining the modes of action of these compounds may be of particular interest for the development of potent inhibitors with therapeutic potential against most NLRP3-AID mutants.

Functional assays on primary patients’ monocytes are limited by the ethical issues related to blood drawn from pediatric patients, and by technical issues related to the low number of monocytes purified from pediatric samples, the cell stress caused by transportation, and the requirement for rapid delivery and processing of clinical samples. Reconstitution of NLRP3-deficient U937 lines with doxycycline-inducible *NLRP3* variants enabled the comparison of *NLRP3* variants on identical genetic background and discriminated NLRP3 constitutive activity *vs* priming-induced and/or activation-induced activity. Noteworthy, expression levels were similar for most of the *NLRP3* variants, excluding any technical artifact caused by overexpression of some mutants. In contrast, the 2 mutants with the lowest expressions, E567G and D303N, were found constitutively active suggesting that their lower expressions likely resulted from their cytotoxic activity. Time-lapse assessment of pyroptosis at the single cell level proved to be a very sensitive and robust readout. Compared to cytokine secretion assays, it required lower numbers of cells and could be applied to monocytes purified from pediatric autoinflammatory patients. IL-18 secretion was poorly reproducible due to its low magnitude and IL-1β was produced only after PMA-differentiation in U937 cell lines. Because PMA-differentiation lowered NLRP3 expression level, and partially provided the priming signal, cytokine secretions appeared to be less robust read-outs. Nevertheless, they confirmed most of the results of pyroptosis assays.

Here we describe five functional groups of *NLRP3* variants, classified according to their cellular response to induction, priming and/or activation signals. This led to the definition of a new group of variants, group#4, responding to the activation signal in the absence of priming. Our data revise previous studies that concluded to the absence of gain-of-function of some of these variants based on unresponsiveness to priming signals despite the higher prevalence in auto-inflammatory patients, and should be considered for atypical patient diagnosis toward NLRP3-AID.^23, 24^ Gain-of-function of most group#4 variants in response to Dox+Nig were of low magnitude, and could be observed using pyroptosis but not cytokine secretion readouts (R168Q, V198M, M701T, Q703K, S726G, Y859C), in line with the incomplete penetrance of these mutations. Noteworthy, some group#5 variants (K375E, G767S, S896P), not considered as gain-of-function based on our analysis, showed trends toward increased pyroptosis in response to Dox+Nig, albeit below the significance threshold. Deeper analysis with more replicates might conclude to slight gain-of-function of these variants. Identification of constitutively active mutants (group#1) by treatment of reconstituted U937 with doxycycline indicated that other key proteins of the pathway (ASC, caspase-1, GSDMD) pre-existed in functional forms in U937, and that priming and activation signals acted on NLRP3 itself.

Given the recent advances in NLRP3 structure determination, the identification of gain-of-function mutations provides insights into the key regulatory mechanisms of inflammasome assembly. A bar diagram depicts the localization of mutant sites according to the domain architecture of NLRP3 (Figure 9A). Based on our current knowledge, the transition from the inactive spherical decamer to the active concentrical decamer requires the release of the LRR domains by binding to NEK7 (Figure 9B). This conformational transition goes along with an 85° rotation of the NBD and HD1 subdomains relative to the WHD and HD2 subdomains within the NACHT, which is the central mechanism of STAND ATPase activation and accompanied by a nucleotide exchange from ADP to ATP (Figures 9C, S13A-B).^14, 25^ Some disease mutations may destabilize the inactive spherical “cage” structure with the PYD effector domains buried inside, that relies mostly on interactions between the LRR domains.^14, 26, 27^ In human NLRP3, a loop section of 42 residues mediates the interlaced LRR dimer assembly with the acidic loop binding on one side into the concave side of its LRR structure and on the other side to the tip of the cognate LRR (Figures 9D, S13C). Y859 maps to the concave side of the LRR directly interacting with the acidic loop (687-700) comprising E690, suggesting that these two NLRP3-AID mutation sites may destabilize the inactive spherical structure.^14^ Residues M701 and Q703 from group#4 similarly reach out into the concave side of the canonical LRR and any mutation here might influence the intertwined LRR dimer assembly.

**Figure 9.**
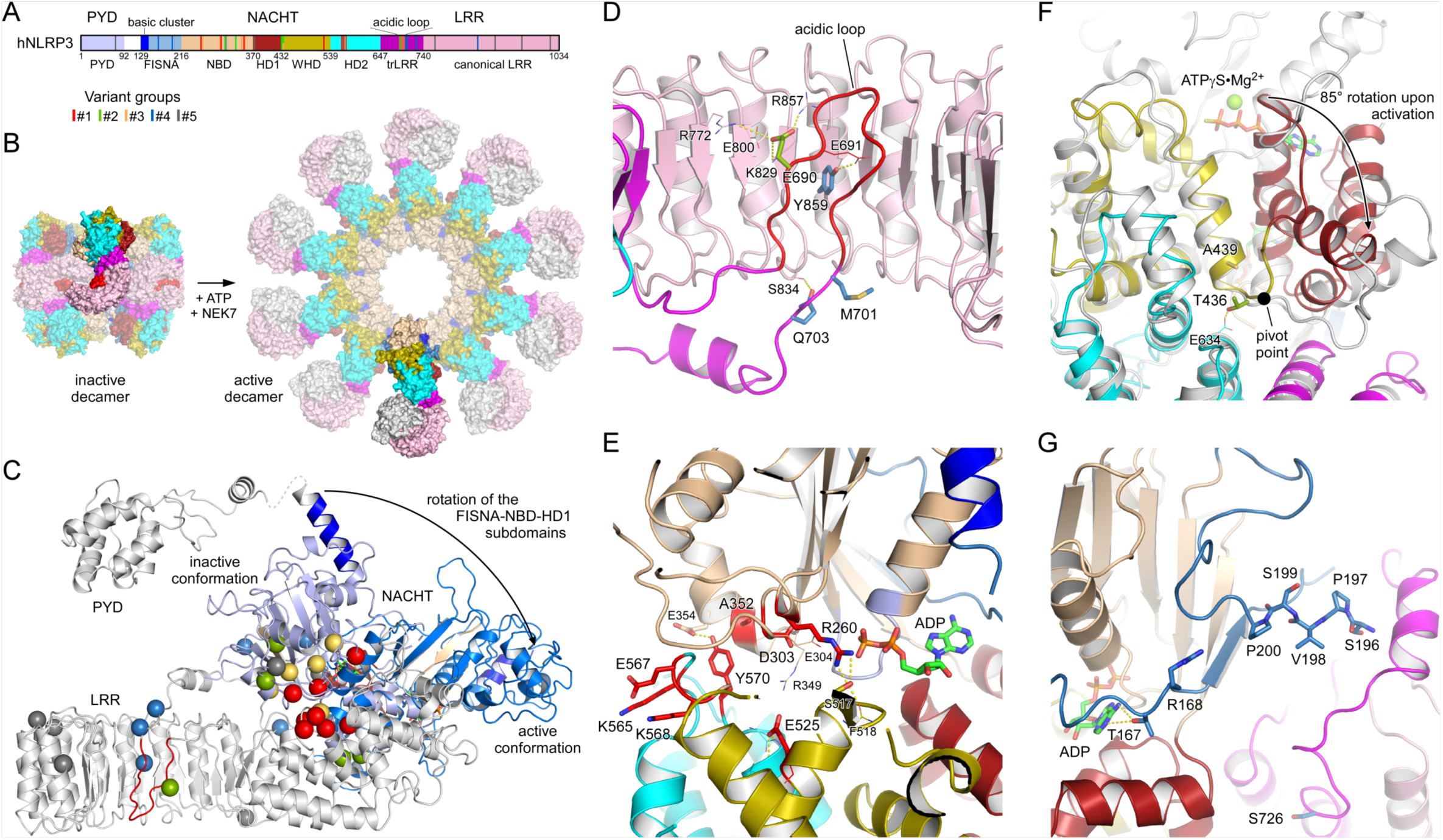
Localization of disease mutant sites in the structure of human NLRP3. A, Bar diagram showing the domain architecture of NLRP3 with the mutant sites of the five variant groups indicated. B, Display of the inactive spherical NLRP3-ADP decamer (7pzc)^12^ and the active concentrical NLRP3-ATP decamer bound to NEK7 (in grey) (8ej4)^25^. C, The conformational transition from the inactive to the active state in NLRP3 involves an 85° rotation of the FISNA-NBD-HD1 subdomains in the NACHT relative to the remainder. The missense mutations of the five variant groups are highlighted as spheres in the inactive conformation. D, Close-up of residues E690, M701 and Q703 in the loop section of the transition LRR and Y859 in the concave side of the LRR. E, Localization of the constitutively active group#1 mutant sites in the NACHT domain of NLRP3. F, Residues T436 and A439 are at the pivot point of the FISNA-NBD-HD1 rotation. The active state (8ej4) is shown in grey relative to the colored inactive state (7pzc). G, Variant group#4 residues R168 and V198 in the FISNA domain and S726 in the transition LRR are located in variable regions of NLRP3.

The group#1 variants of constitutively active mutants locate all in the NACHT domain in proximity to the nucleotide binding site. D303 is in the Walker B motif with the typical aaaaDE motif degenerated to aaaaDGxDE in NLRs (where “a” denotes an aliphatic residue).^15^ The two acidic residues catalyze the intrinsic ATP hydrolysis mechanism that is required for the relapse from an active to an inactive state, and it is well perceived that any mutation in this motif enriches the active state of NLRP3. A352 is following the sensor 1 motif (R349) on β-strand 4, interacting with D303 on β-strand 3 from the back (Figure 9E). Its mutation to a larger residue could also disrupt the inactivation mechanism. The R260 residue at the tip of β-strand 2 is buried in the structure of the inactive NACHT domain (7pzc) but on the surface close to the γ-phosphate group in the active NACHT conformation (8ej4). Its mutation to a larger Trp-residue might prevent the formation of the tightly closed inactive state. The acidic E525 in WHD forms a salt bridge with the sensor 1 R349 following H-bonds with HD2. A charge reversal to Lys will dismiss these interactions. Finally, residues K565, E567 and K568 are on the accessible surface of the HD2 domain, both in the inactive and active state. Their disease related mutations point to an important surface patch of this side. The succeeding residue Y570 is again inside and interacts with the NBD, yet its mutation to Cys could indirectly influence the three preceding residues. This similarly holds for the G569R mutation of group#3, which might also affect the integrity of the K_565_xEKGY stretch.

The group#3 G301D mutant is proximate to D303 and part of the degenerated Walker B motif. The G301E mutation has been shown to have an eight-fold reduced hydrolysis activity, enriching thus the active state. Residues T436 and A439 from groups #2-3 are located in the WHD near the pivot point switching between the inactive and active conformation (Figure 9F).^14, 25^ The new group#4 disease variants R168 and V198 are both located in the FISNA domain. R168 is next to T167 that interacts with the adenine base of the bound nucleotide (Figure 9G). V198M may stabilize the FISNA domain in the active conformation, as V198 is located in the activation loop comprising S_196_PVSP, whose priming-associated phosphorylation is likely to destabilizes the inactive cage complex.^14, 25^

MCC950 binds NLRP3 in a cleft spanned by subdomains NBD, HD1, WHD, HD2 and transition LRR, stabilizing the inactive conformation of the spherical decamer (Figure S13D).^14^ Our results on the inhibition of cell death upon NLRP3 WT or NLRP3-AID variant expressions confirmed the sensitivity to MCC950 of V198M, E311K, T348M, the partial sensitivity of A352V, A439V, D303N, and the full resistance of L353P.^12, 16, 20, 28^ D303H and E525K (group#1) and G301D, L353P and G569R (group#3) were fully resistant to MCC950, either because they do not bind to MCC950, such as L353P,^20^ or MCC950 is not sufficient to stabilize the inactive conformation. The group#3 mutants were activated upon priming, indicating that the mechanism of NLRP3 priming may not rely solely, or at all, on an inactive cage structure destabilization. Indeed, mutants activated by the activation signal nigericin without priming (group#4) were highly sensitive to MCC950 to a similar extent as the WT NLRP3, suggesting that these mutants form inactive structures and that the activation signal is at least involved in destabilizing this inactive form. Alternatively, MCC950 may inhibit NLRP3 activity independently of stabilizing the cage *per se*, but rather the closed conformation of the NACHT domain, e.g. when bound to NEK7, or additional other mechanisms.

While CY-09 is also considered a direct inhibitor of NLRP3, its action mechanism is poorly characterized.^13^ It is supposed to compete with nucleotide-binding to NLRP3; yet, a recent study has reported that CY-09 does not inhibit NLRP3 ATPase activity.^15^ Our results partly confirmed NLRP3 A352V (group#1) sensitivity to CY-09.^13^ In contrast to MCC950, no gain-of-function mutations showed full resistance to CY-09, consistent with CY-09 targeting residues indispensable for NLRP3 inflammasome assembly, suggesting that CY-09 and its derivatives may have broader activity than MCC950 for future clinical perspectives against NLRP3-AID. However, as a proposed ATP-competitive inhibitor, CY-09 might exhibit many off-target effects.

G5 is a deubiquitinase inhibitor that maintains NLRP3 in an ubiquitinated inactive form.^18^ Consistent with the lack of specificity of G5 targeting multiple deubiquitinases, and the reports of multiple ubiquitination sites on NLRP3, none of the *NLRP3* gain-of-function mutants showed resistance to G5.^18, 29–31^ Among all mutants, NLRP3 A352V (group#1) and T436N (group#2) showed lower sensitivity to G5 upon treatment with doxycycline only and LPS only, respectively. However, as the amplitude of the activation was low in these conditions, the low percentage of inhibition may be artefactual and should be interpreted with caution. Noteworthy, constitutively active mutants were inhibited by G5, indicating that ubiquitination and deubiquitination may occur at least in part independently of priming and activation signals. The hypothesis that G5 may have an additional target downstream of NLRP3 in the pathway is ruled out by the specificity of G5 to inhibit the NLRP3 but neither the AIM2 nor the NLRC4 inflammasomes.^18^

CRT0066101 is a pan-PKD inhibitor targeting PKDs-mediated NLRP3 S293 phosphorylation upon activation signals that controls the release of NLRP3 from an intracellular membrane compartment to the cytosol for the assembly of the inflammasome.^19^ None of the group#2 and #4 mutants were inhibited by CRT0066101 when activated by activation signal in the absence of priming (Figure 7, Dox+Nig only), indicating that in this context NLRP3 activation occurs independently of PKDs. Further investigation would be necessary to test if these mutants are released from the intracellular membranes independently of PKDs, display activity while associated to the membranes, or if they are not recruited to the membranes. In the latter hypothesis, priming might control NLRP3 recruitment to intracellular membranes or may disrupt an alternative pathway that bypasses the requirement for PKD by activating another kinase or another mechanism to control NLRP3 release from the intracellular membrane. Constitutive activity of most of the group#1 mutants were sensitive to CRT0066101, indicating that basal PKD activity may be required.

Altogether, our study constitutes the most comprehensive comparative analysis of *NLRP3* mutations associated with auto-inflammation reported so far. Our results reveal the functional diversity of *NLRP3* gain-of-function mutations that should be taken in consideration for atypical NLRP3-AID diagnosis and specific targeted treatments perspectives.

## Methods

### Study approval

The study was carried out in the setting of the ENFLAMAI protocol, which was previously approved by the French *Comité de Protection des Personnes* (CPP, #L16-189) and by the French *Comité Consultatif sur le Traitement de l’Information en matière de Recherche dans le domaine de la Santé* (CCTIRS, #16.864). The experiments conformed to the principles set out in the WMA Declaration of Helsinki and the Department of Health and Human Services’ Belmont Report. Anonymous healthy donors’ blood was provided by the *Etablissement Français du Sang* (EFS) in the framework of the convention #14-1820 between Inserm and EFS. Informed consent was received from participants prior to inclusion in the study.

### NLRP3 variants nomenclature

All NLRP3 amino-acids are numbered according to the Infevers database European nomenclature (https://infevers.umai-montpellier.fr/web/).^32^ Their pathogenicity was classified by an International consortium of expert geneticists as previously described.^4^

### Cell culture

Lenti-X^TM^ 293T cells (Takara) were cultured at 0.5-1x10^6^ cells/ml in Dulbecco’s modified Eagle’s Medium (DMEM) GlutaMax^TM^-I supplemented with 1X penicillin/streptomycin (PS) and 10% Fetal Bovine Serum (FBS) (Gibco). NLRP3-deficient U937 cells have been previously described. U937 and U937 reconstituted NLRP3-deficient cells were cultured at 0.25-1x10^6^ cells/ml in Roswell Park Memorial Institute (RPMI) 1640 GlutaMax^TM^-I supplemented with 1X PS and 10% FBS (Gibco). Monocytes were purified from 10-20 ml of blood samples (drawn the day before in heparin tubes) using EasySep^TM^ Direct Human Monocyte Isolation kit (StemCell) and cultured at 0.07-0.15x10^6^ cells/ml in RPMI-1640 GlutaMax^TM^-I supplemented with 1X PS and 10% FBS (Gibco).

### Plasmids

Human *NLRP3* were cloned in pENTR^TM^1A (Invitrogen) from Flag-NLRP3 encoding plasmids kindly shared by F. Martinon (Unil, Lausanne, Switzerland). Mutations were performed using QuickChange^TM^ II kit (Agilent Technologies). cDNAs were transferred in pInducer21 using recombination Gateway LR clonase Enzyme mix kit (Thermofisher).^33^ pMD2.G and pCMVR8.74 were gifts from D. Trono (Addgene plasmid #12259 and #22036).

### Reagents

The following reagents were used: polyethylenimine (PEI, Sigma), polybrene (Santa-Cruz), phorbol myristate acetate (PMA, Invivogen), doxycycline (Sigma), LPS (O111:B4, Sigma), nigericin (Invivogen), ATP (Sigma), MCC950 (Adipogen), CY-09 (tocris), G5 (Calbiochem), CRT0066101 (Tocris), propidium iodide (PI, Immunochemistry technologies), Hoechst (Immunochemistry technologies)

### Lentivector preparation and cell transduction

1.6x10^6^ Lenti-X^TM^ 293T cells were co-transfected with pMD2.G (10 mg), pCMVR8.74 (30 mg) and pInducer21-NLRP3 or its variants (40 mg) using PEI to produce lentivectors. Lentivectors were concentrated by ultracentrifugation (30,000 rpm, 1h) and used to transduced 1.5x10^5^ NLRP3-deficient U937 cells in the presence of polybrene (8mg/ml)^11^. 4 days later, transduced GFP-positive cells were sorted by cytometry as bulk.

### Western-blot analysis

To assess NLRP3 protein levels, U937 cells were differentiated with PMA (50 ng/ml, 16h) and treated with LPS (40 ng/ml, 3h) or doxycycline (0.5-2 mg/ml, 3h). Cells were directly lysed in sample buffer 2X. Samples were analyzed by SDS-PAGE and transferred to PVDF membranes. The following antibodies were used: anti-NLRP3 (Cryo2, Adipogen), anti-IL-1β (D3U3E, Cell Signaling), anti-IL-18 (D2F3B, Cell Signaling), anti-LaminB (B-10, Santa Cruz Biotechnology), anti-actin (C4, Sigma), HRP-anti-Mouse IgG (H+L) (Promega).

### Cell death assay

To assess U937 permeabilization, 0.3×10^6^ cells/ml were platted in media without phenol red. The next day, cells were treated with doxycycline (1 μg/ml, 3h), LPS (40 ng/ml, 2h) and nigericin (15 μg/ml) before time-lapse imaging using CQ1 high content screening microscope (Yokogawa). In inhibition assay, MCC950 (1 μM), CY-09 (50 μM), G5 (1 μM), CRT0066101 (0.5 μM) or DMSO vehicle (1% final in all conditions) were added 20 min before doxycycline, LPS or nigericin as indicated. To assess monocyte permeabilization, 0.07-0.15×10^6^ cells/ml were platted in media without phenol red and directly treated with MCC950 (1 μM, 3h15), LPS (40 ng/ml, 3h) and nigericin (5 μg/ml) before time-lapse imaging using CQ1 high content screening microscope (Yokogawa). PI (1.25 μg/ml) and Hoechst (0.2 μg/ml) were added 2h before imaging. 2 images/well were taken every 15 min for 2h using 10X objectives (UPLSAPO 10X/0.4). Images were quantified using the CQ1 software (Yokogawa).

### Cytokine secretion assay

To assess IL-18, IL-1β and TNF secretion, 0.25×10^6^ reconstituted U937 cells/ml were differentiated with PMA (50 ng/ml, 16h). The next day, cells were treated with doxycycline (1 μg/ml, 4h), LPS (40 ng/ml, 3h) and nigericin (15 μg/ml, 1h). Cell-free media were then analyzed using IL-18, IL-1β and TNF ELISA kits (RnD Systems).

### Statistical analysis

The cell death was observed at 10 time points ranging from 0 to 135 minutes by 15 minutes interval for each of the 34 *NLRP3* variants and *NLRP3* WT undergoing 5 treatments (*i.e.* untreated, Dox, Dox+LPS, Dox+Nig and Dox+LPS+Nig), and studied as a proportion of the whole cell population.

The database was divided into 50 data subsets (5 treatments * 10 time points). For each subset we fitted a generalized linear mixed model (glmm), beta family with logit link producing 50 results for each variant. In our model, the *NLRP3* variant type was set as the explanatory variable with the WT as the reference. The outcome variable in the model was the frequency of dead cells bounded in [0,1]. glmm can similarly to lmm correct for the technical batch effect induced by the individual experiment. The p-values obtained were corrected by the Bonferroni approach to take into account results for the 5 treatments. We used the R statistical environment, version 4.1.3, with the glmmTMB function from the glmmTMB package. Results for Dox, Dox+LPS and Dox+Nig treatments are shown. For visualisation purpose, cut-offs were fixed to regression coefficients 1.5/-1.5 for Dox and Dox+LPS, and 0.5/-0.5 for Dox+Nig, with p-value<0.05. As data examples, raw data of duplicates of representative experiments were analyzed by two-way ANOVA multiple comparison using *NLRP3* variant and treatments as variable (Graphpad, Prism). Experiments were independently repeated 2-8 times.

IL-1β, IL-18 and TNF concentrations were measured for the 34 *NLRP3* variants and *NLRP3* WT upon 3 treatments (*i.e.* Dox, Dox+LPS, Dox+Nig). One dataset per treatment was created, and a linear mixed-effects model (lmm) was realized for each dataset to assess the dependence of the cytokine concentration on the variant. Each dataset comprised 564 observations. The outcome in the model was the log1p (log1p(X)=log(1+X)) of the cytokine concentration given in pg/ml while the right side of the model was the *NLRP3* variant. The *NLRP3* WT was set as the reference. lmm was appropriate due to its ability to correct for the batch effect triggered by the individual experiment. We used the R statistical environment, version 4.1.3, with the lme function from the nlme package. For visualisation purpose, cut-offs were fixed to regression coefficients 1/-1 with p-value<0.05. As data examples, raw data of duplicates of representative experiments were analyzed by two-way ANOVA multiple comparison using *NLRP3* variant and treatments as variable (Graphpad, Prism). Experiments were independently repeated 3-18 times.

The effect of the NLRP3 inhibitors was tested by comparing AUC obtained in presence of each of the 4 inhibitors (*i.e.* MCC950, CY-09, G5 and CRT0066101) to the AUC obtained without inhibitor (vehicle only, used as the reference state). A linear mixed-effects model (lmm) was fitted for each genotype undergoing a given treatment, in a dataset constructed by the extraction of genotype*treatment from all data. The model response variable was log10(AUC) and the explanatory variable was the inhibitors. As data diagnosis proved a lawful steady batch effect of the independent experiment, lmm modeling was used to correct for this batch effect technical bias in the data. The p-values obtained were corrected by the Bonferroni approach to take into account results for the 4 treatments (Dox, Dox+LPS, Dox+Nig and Dox+LPS+Nig). For visualisation purpose, cut-offs were fixed to regression coefficients <-0.1 with p-value<0.05. Regression coefficient=log10(Fold change). We used the R statistical environment, version 4.1.3, with the lme function from the nlme package. As data examples, AUC of duplicates of representative experiments were analyzed by two-way ANOVA multiple comparison using treatments and inhibitors as variable (Graphpad, Prism). Experiments were independently repeated twice.

## Acknowledgements

This work was supported in part by the the Institut National de la Santé et de la Recherche Médicale (Inserm), the Centre National de la Recherche scientifique (CNRS), ENS Lyon, Université Claude Bernard Lyon 1 (UCBL1), the European Research Council (ERC-2013-CoG_616986 to BP), the Agence Nationale de la Recherche (ANR-13-JSV3-0002-01 and ANR-22-CE15-0032-01 to BP, ANR-18-CE17-0001 “Action” and ANR-18-CE15-0017) and through the “Investissements d’avenir” program (Institut Hospitalo-Universitaire Imagine, ANR-10-IAHU-01; Recherche Hospitalo-Universitaire, ANR-18-RHUS-0010), the Fondation pour la Recherche Médicale (Equipe FRM EQU202103012670 to FRL). We thank Fabio Martinon (UNIL, Lausanne, Switzerland) for the FLAG-NLRP3 constructs, and Didier Trono for the pMD2.G and pCMVR8.74 (EPFL, Lausanne, Switzerland). We thank Isabelle Durieux, Hermine Achard and Marie Guirguis (CIRI, Lyon, France) for help with subcloning. We acknowledge the contribution of SFR Biosciences (UMS3444/CNRS, US8/Inserm, ENS de Lyon, UCBL) PLATIM and flow cytometry facilities, as well as of the Etablissement Français du Sang Auvergne - Rhône-Alpes.

## Conflict of Interest statement

The authors declare no conflicting financial interests.

## Supplementary Information

**Supplemental Figure 1.**
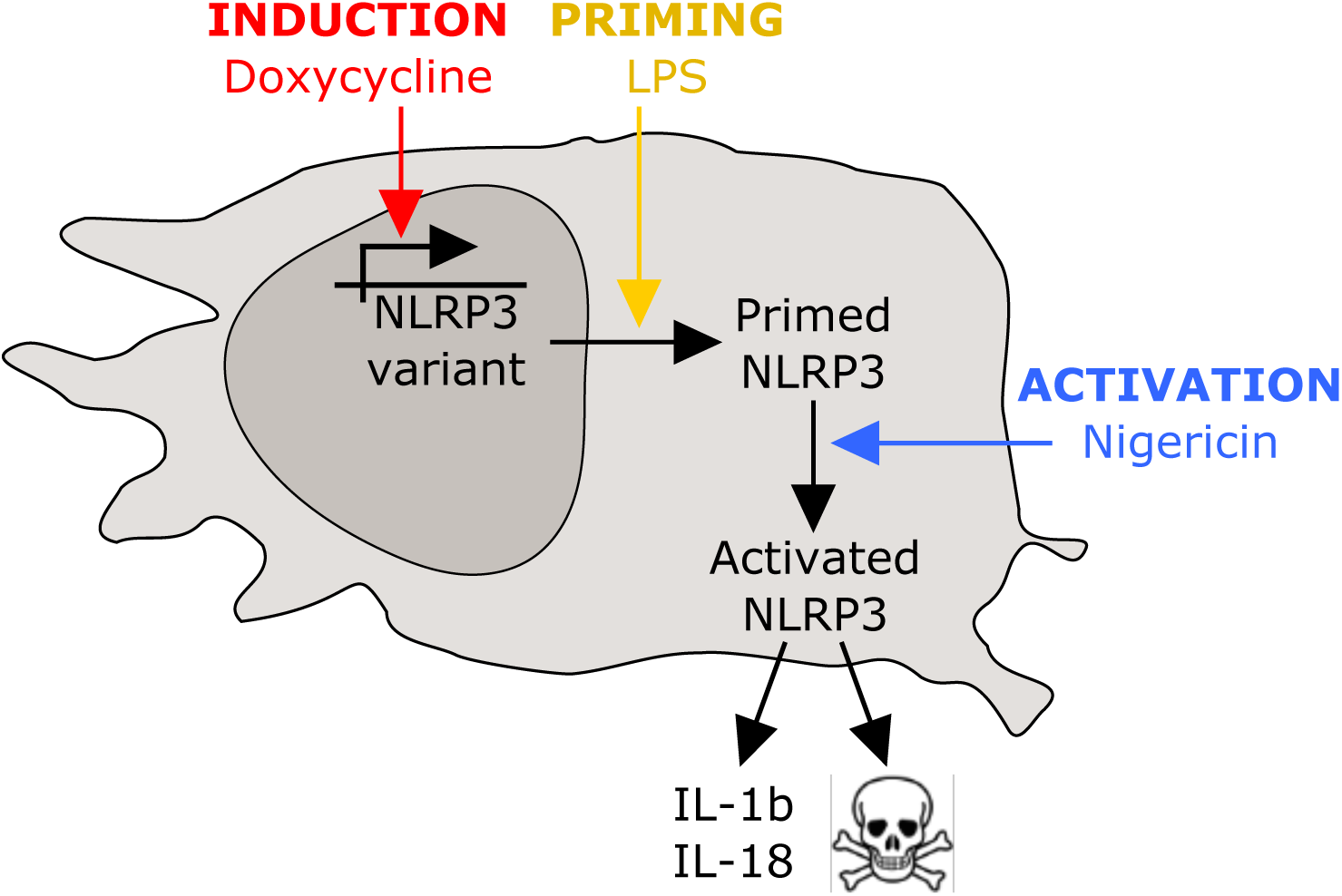
Assessment of the activity of *NLRP3* variants upon expression, priming and/or activation signals. *NLRP3*-deficient KO U937 reconstituted with doxycycline-inducible *NLRP3* variants were treated with doxycycline, LPS priming signal and/or nigericin activation signal. NLRP3 activity was assessed by measuring pyroptosis and IL-1β/18 secretion.

**Supplemental Figure 2.**
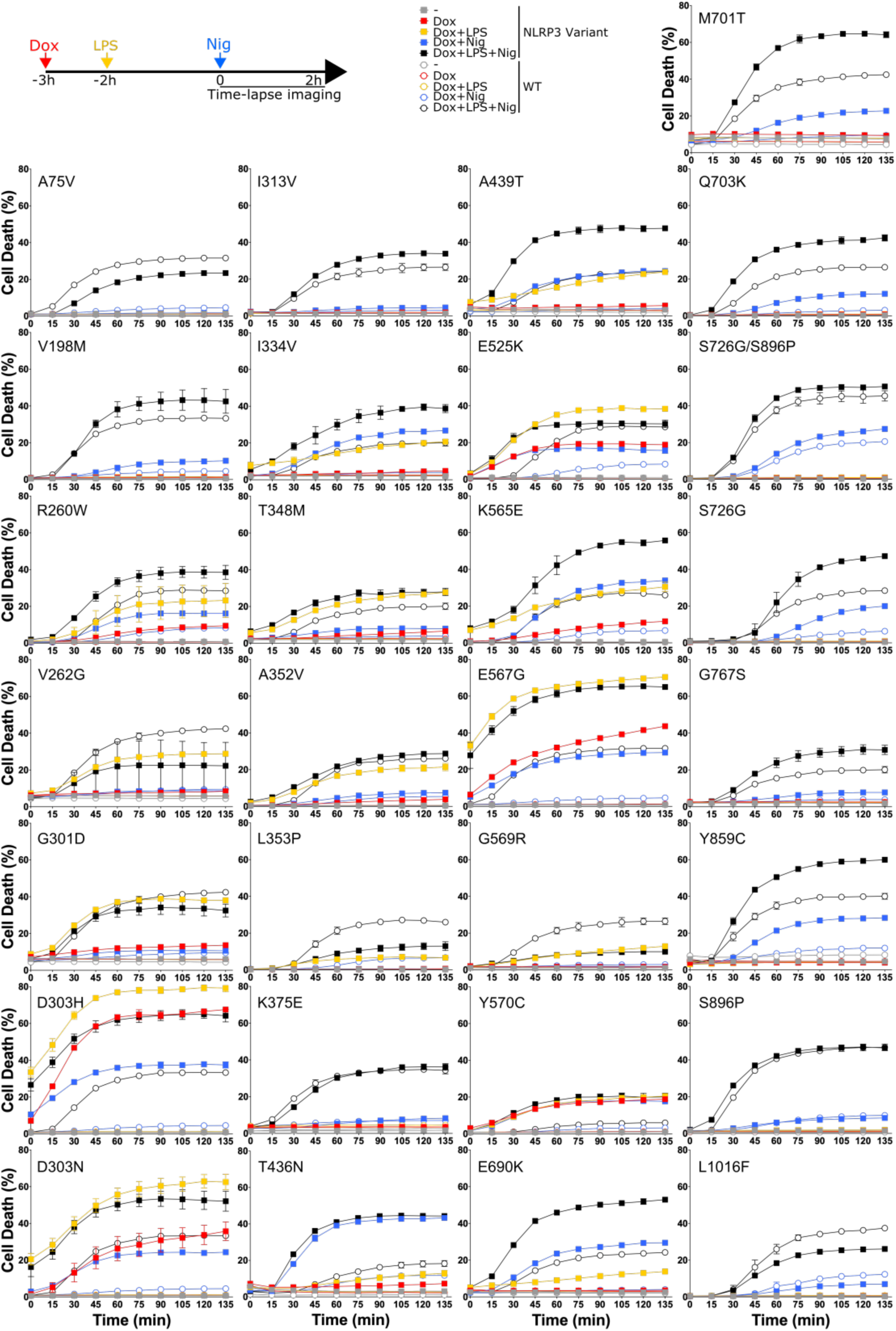
NLRP3 expression, priming and/or activation triggers pyroptosis in reconstituted U937 depending on the *NLRP3* variants. *NLRP3*-deficient U937 cells reconstituted with doxycycline-inducible *NLRP3* variants were treated with doxycycline (1 μg/ml, 3h), LPS (40 ng/ml, 2h) and nigericin (15 μg/ml) before cell death was monitored by PI incorporation over time quantified by time-lapse high content microscopy. Means of duplicates and 1 SD are represented. One experiment done in duplicates representative of 2-8 independent experiments is shown. Statistical analysis including all independent experiments are represented in Figure 3.

**Supplemental Figure 3.**
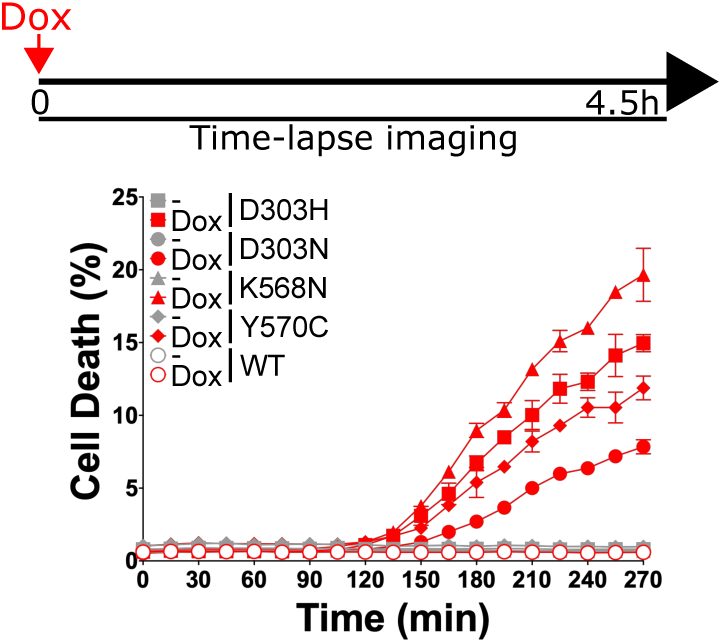
Kinetics of doxycycline-induced death of U937 expressing group#1 *NLRP3* variants. *NLRP3*-deficient U937 cells reconstituted with doxycycline-inducible group#1 *NLRP3* variants were treated with doxycycline (1 μg/ml) before cell death was monitored by PI incorporation over time quantified by time-lapse high content microscopy. One experiment done in duplicates representative of 2 independent experiments is shown.

**Supplemental Figure 4.**
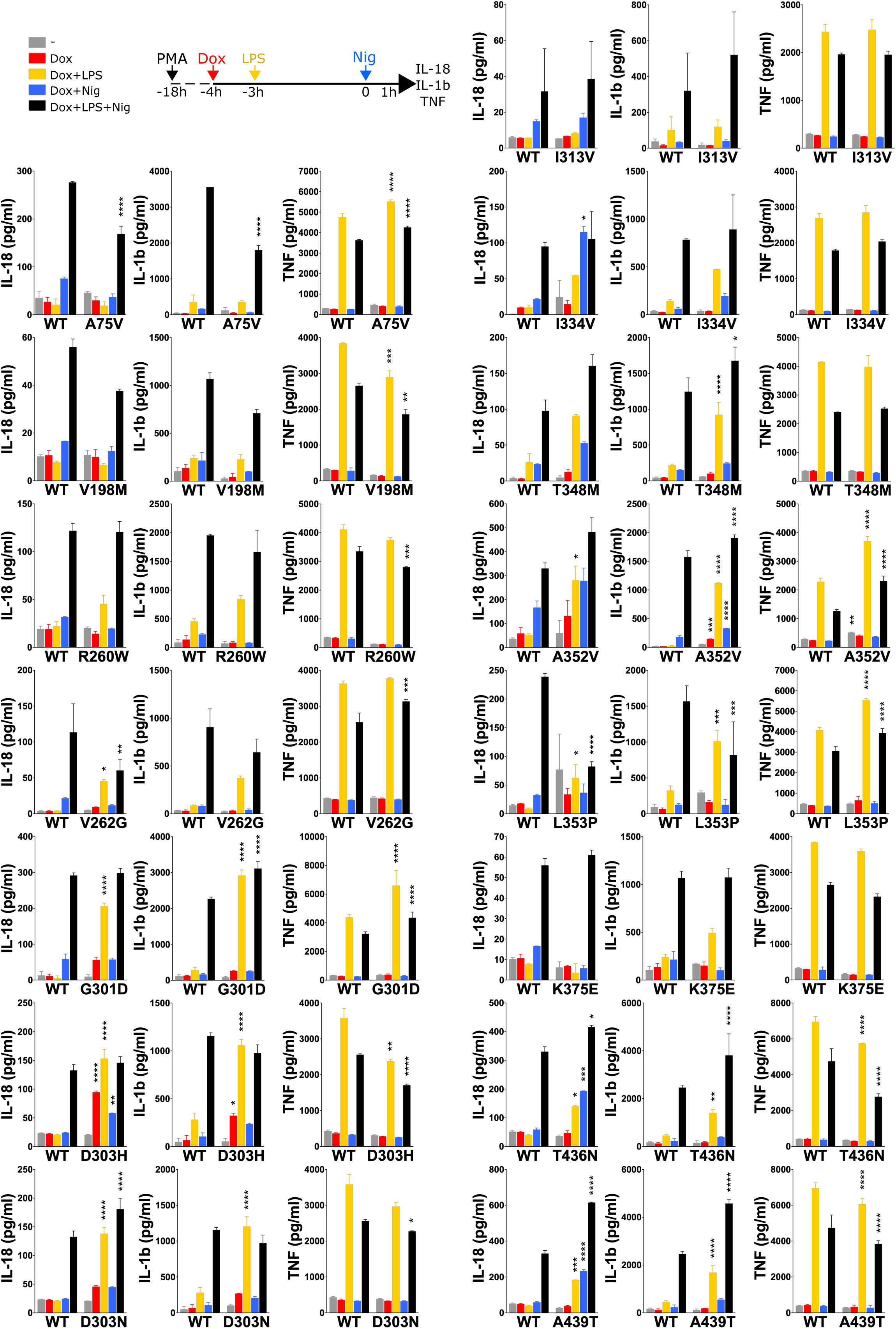
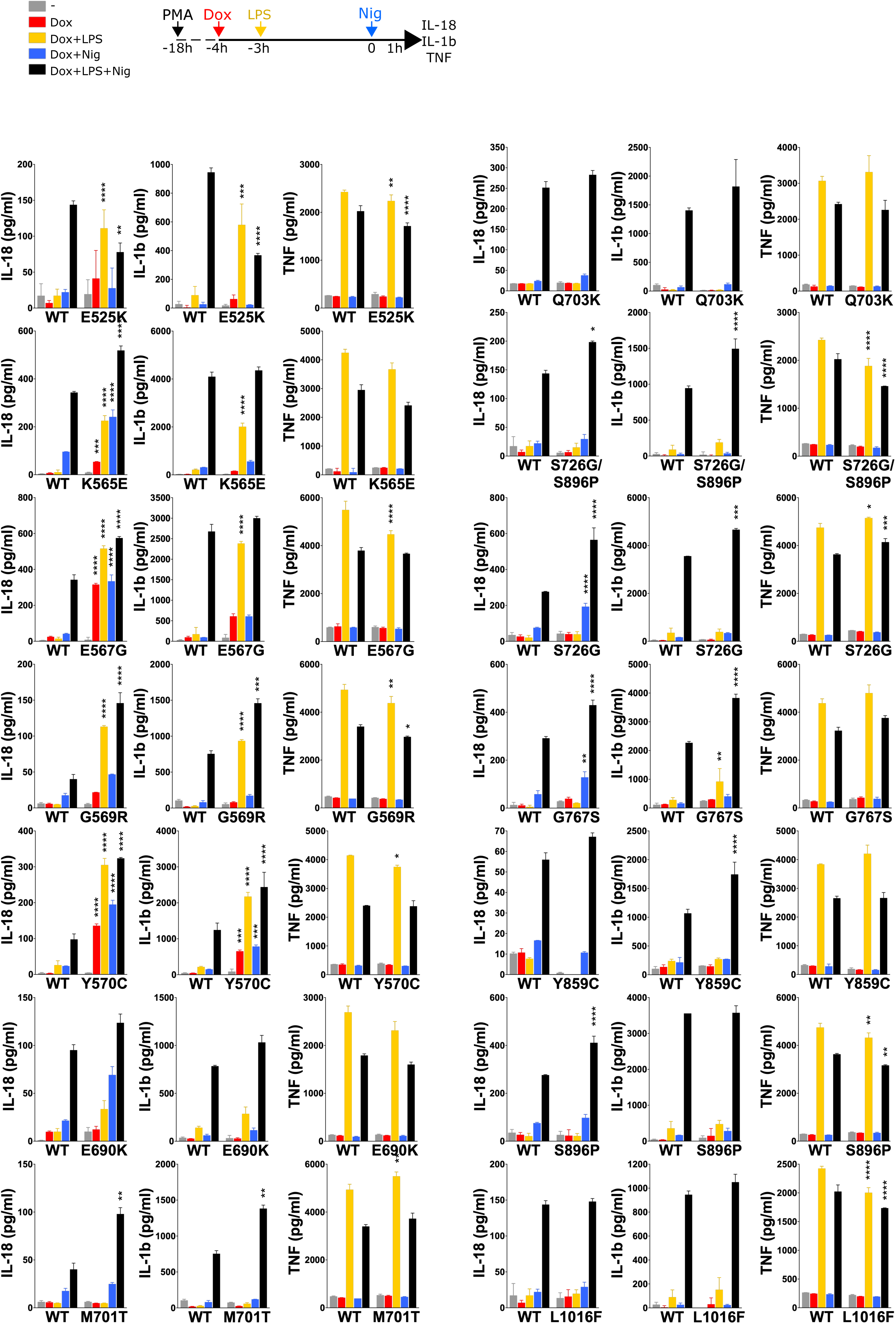
NLRP3 expression, priming and/or activation triggers IL-18 and IL-1β secretion in reconstituted U937 depending on the *NLRP3* variants. *NLRP3*-deficient U937 cells reconstituted with doxycycline-inducible *NLRP3* variants were differentiated in PMA (50 ng/ml) and treated with doxycycline (1 μg/ml), LPS (40 ng/ml) and nigericin (15 μg/ml) as indicated. IL-18, IL-1β and TNF (as control) levels were assessed in cell supernatant by ELISA. Means of duplicates and 1 SD are represented. Two-way ANOVA multiple comparisons of each variant with WT control with corresponding treatment on the same plate, *, p <0.05; **, p <0.01; ***, p <0.001; ****, p <0.0001. One experiment done in duplicates representative of 3-18 independent experiments is shown. Statistical analysis including all independent experiments are represented in Figures 5 and S5.

**Supplemental Figure 5.**
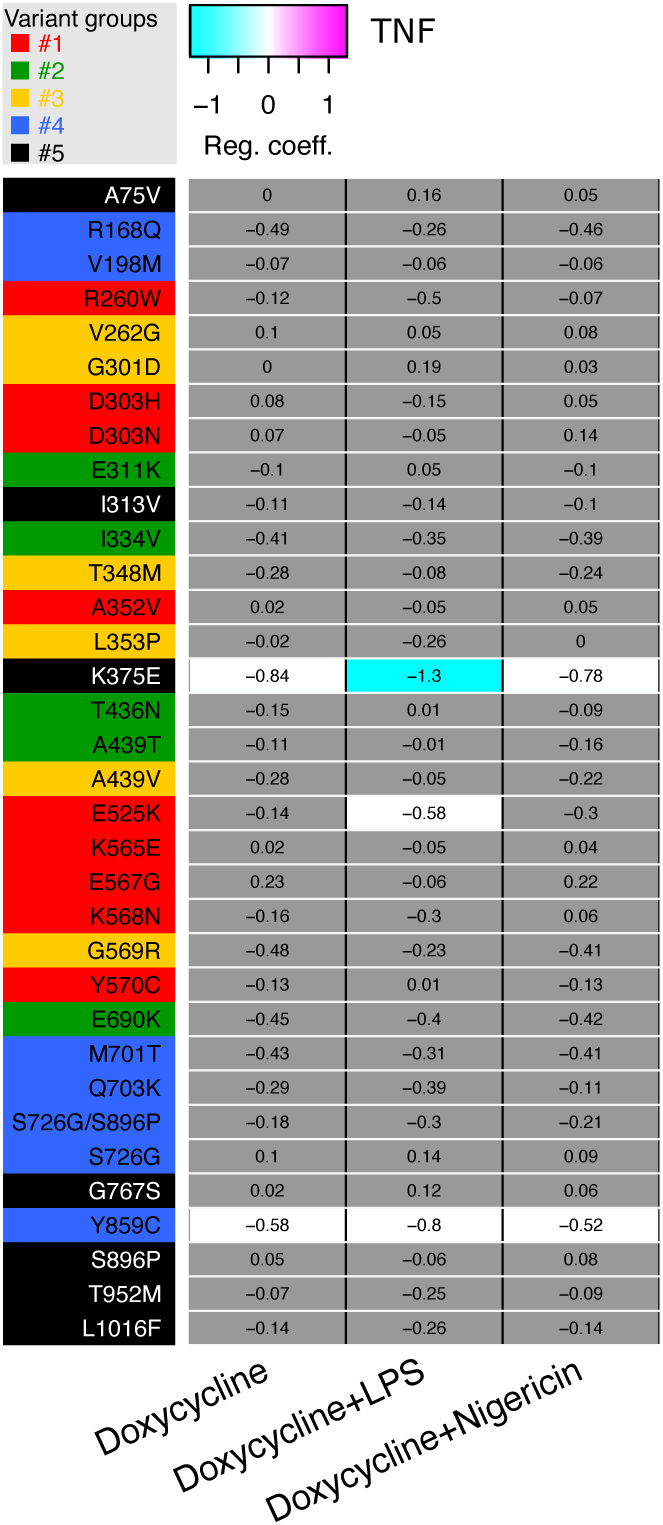
Relative TNF secretion of U937 expressing *NLRP3* variants as compared to U937 expressing WT *NLRP3* in response to NLRP3 expression, priming and/or activation. Regression coefficient (RC) heatmap from the lmm modeling for each variant as compared to the WT. Positive RC denotes increased TNF concentrations by the variant as compared to *NLRP3* WT, and conversely. RC corresponding to statistically significant increases (magenta) or decreases (cyan) in cytokine secretions are color-coded (p<0.05 and, RC>+1 or RC<-1 respectively). Not significant, p>0.05 (grey).

**Supplemental Figure 6.**
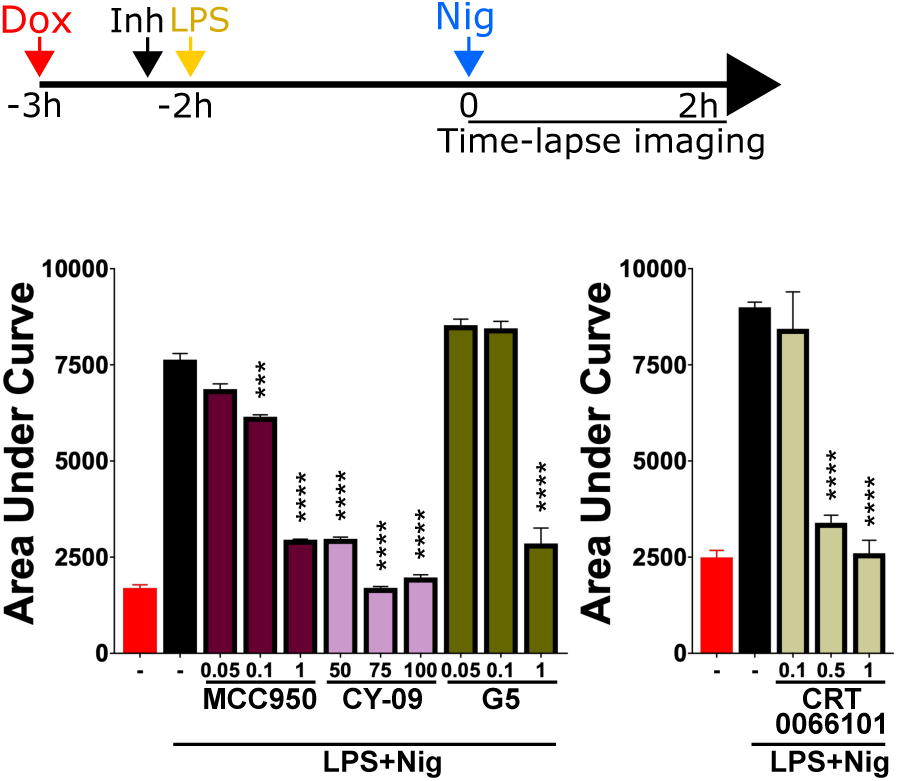
Dose-dependent inhibition of pyroptosis by NLRP3 inhibitors. *NLRP3*-deficient U937 cells reconstituted with doxycycline-inducible WT *NLRP3* and treated with doxycycline (1 μg/ml, 3h) were treated with indicated doses (μM) of MCC950, CY-09, G5 and CRT0066101 or vehicle (2h15), LPS (40 ng/ml, 2h) and nigericin (15 μg/ml) before cell death was monitored by PI incorporation over time quantified by time-lapse high content microscopy. Means of area under the curve of duplicates and 1SD are represented. Two-way ANOVA multiple comparisons of each variant with WT control with corresponding treatment, ***, p <0.001; ****, p <0.0001.

**Supplemental Figure 7.**
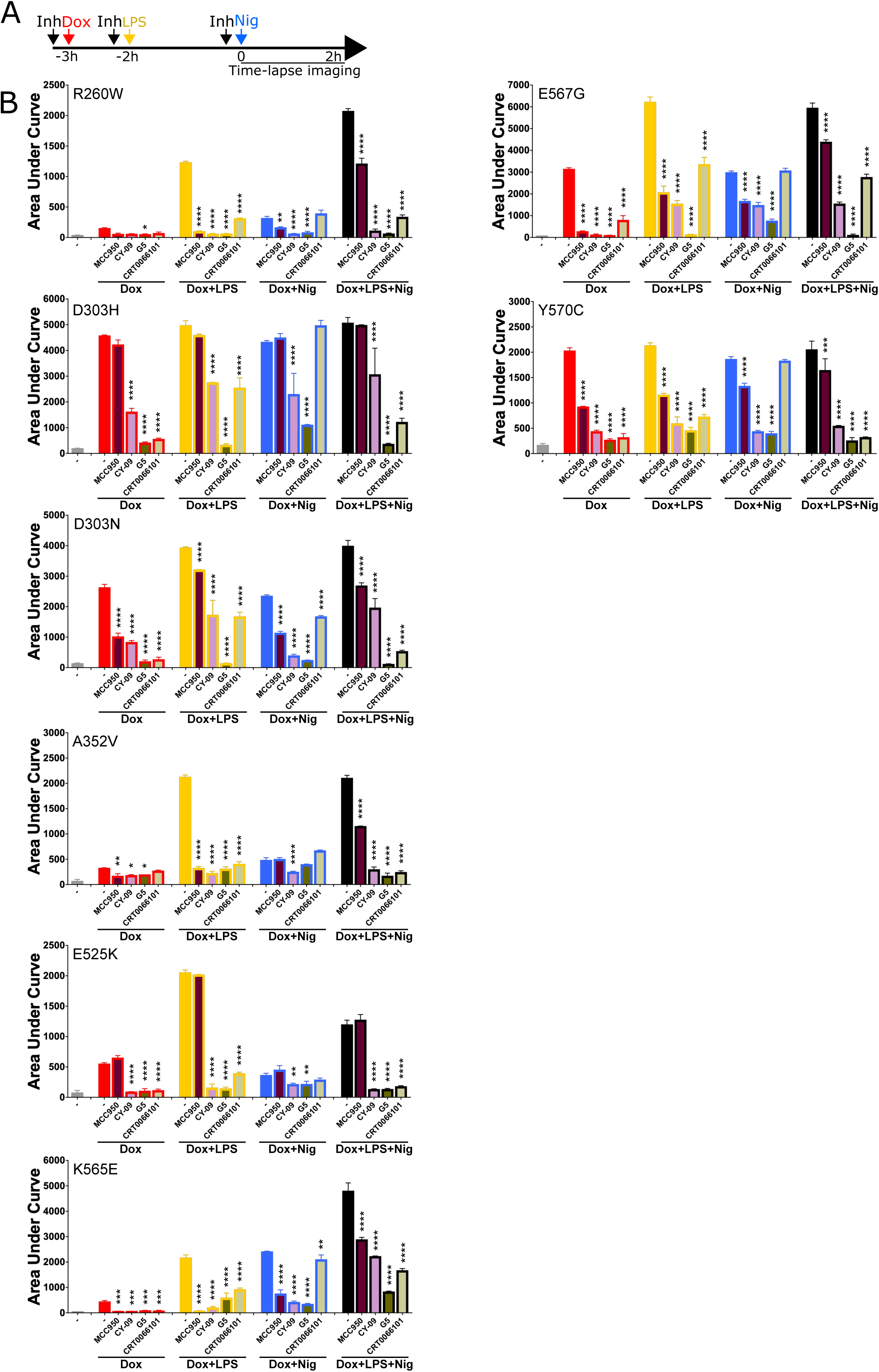
Sensitivity of group#1 *NLRP3* variants to NLRP3 inhibitors. A-B. *NLRP3*-deficient U937 cells reconstituted with doxycycline-inducible *NLRP3* group#1 variants were treated with doxycycline (1 μg/ml, 3h), LPS (40 ng/ml, 2h) and/or nigericin (15 μg/ml) in the presence of MCC950 (1 μM), CY-09 (50 μM), G5 (1 μM) and CRT006101 (0.5 μM) NLRP3 inhibitors or DMSO vehicle (added 20 min before the last treatment), and cell death was monitored by PI incorporation over time quantified by time-lapse high content microscopy for 2h. Means of area under the curve of duplicates and 1SD are represented. Two-way ANOVA multiple comparisons of each inhibitor with the corresponding treatment with vehicle only. *, p <0.05; **, p <0.01; ***, p <0.001; ****, p <0.0001. One experiment done in duplicates representative of 2 independent experiments is shown. Statistical analysis including all independent experiments are represented in Figure 7.

**Supplemental Figure 8.**
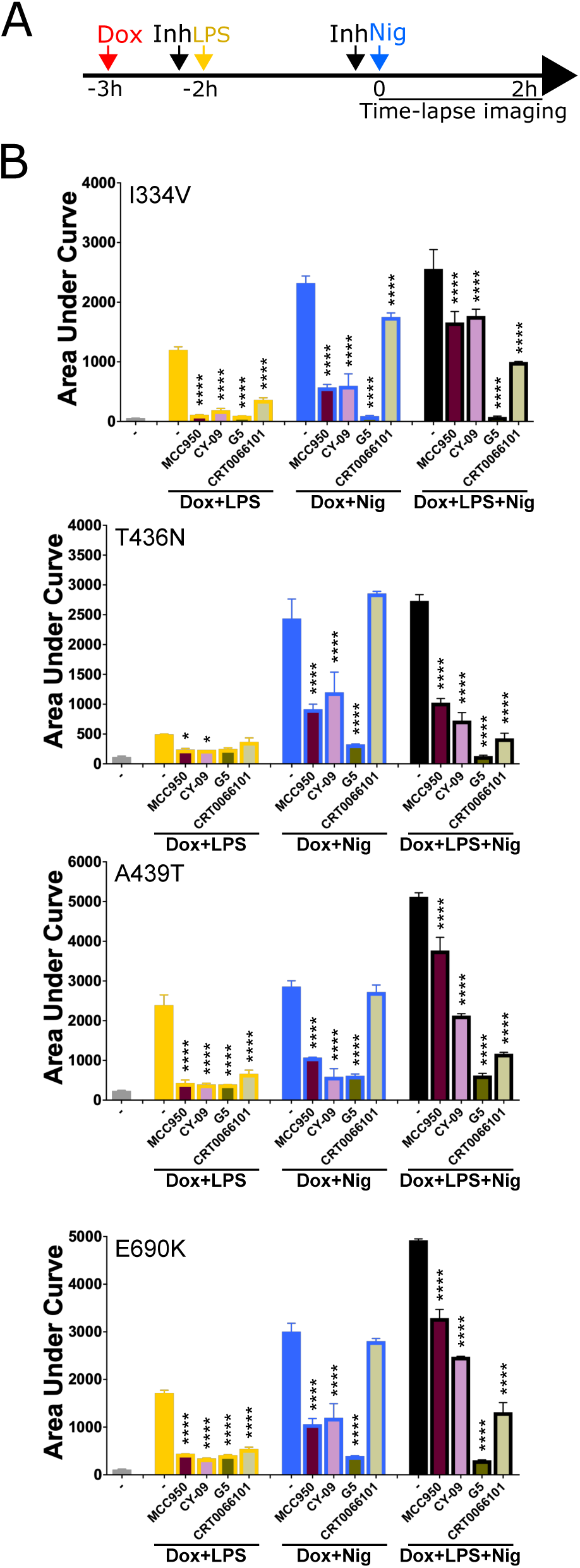
Sensitivity of group#2 *NLRP3* variants to NLRP3 inhibitors. A-B. *NLRP3*-deficient U937 cells reconstituted with doxycycline-inducible *NLRP3* group#2 variants were treated with doxycycline (1 μg/ml, 3h), LPS (40 ng/ml, 2h) and /or nigericin (15 μg/ml) in the presence of MCC950 (1 μM), CY-09 (50 μM), G5 (1 μM) and CRT006101 (0.5 μM) NLRP3 inhibitors or DMSO vehicle (added 20 min before the last treatment), and cell death was monitored by PI incorporation over time quantified by time-lapse high content microscopy for 2h. Means of area under the curve of duplicates and 1SD are represented. Two-way ANOVA multiple comparisons of each inhibitor with the corresponding treatment with vehicle only. *, p <0.05; **, p <0.01; ***, p <0.001; ****, p <0.0001. One experiment done in duplicates representative of 2 independent experiments is shown. Statistical analysis including all independent experiments are represented in Figure 7.

**Supplemental Figure 9.**
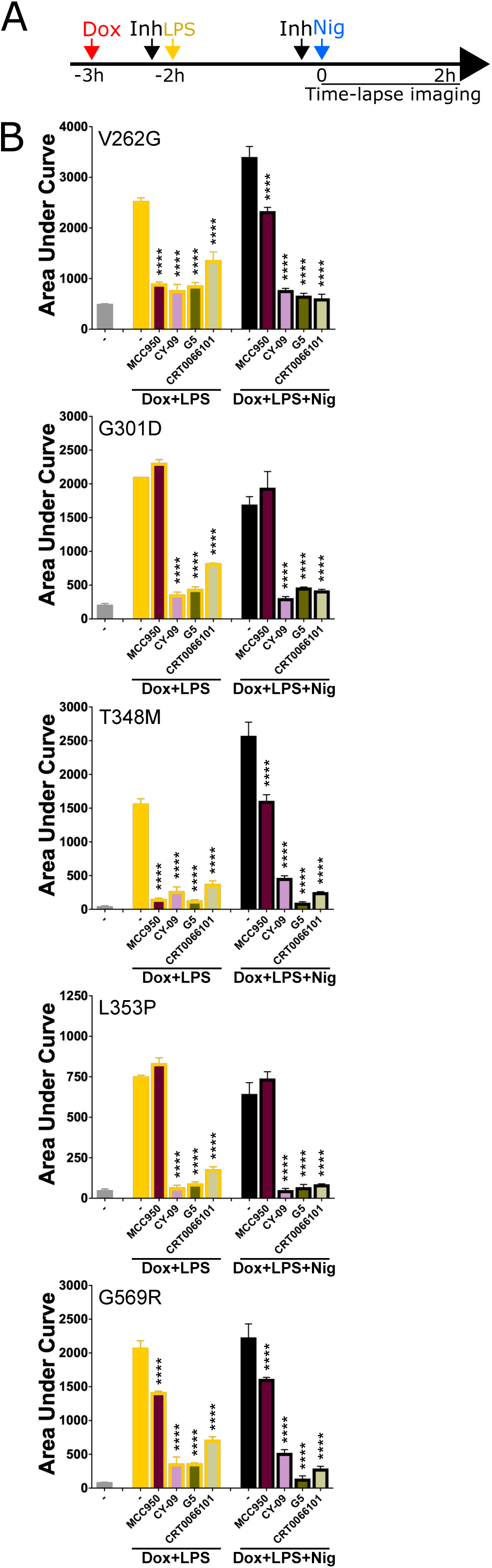
Sensitivity of group#3 *NLRP3* variants to NLRP3 inhibitors. A-B. NLRP3-deficient U937 cells reconstituted with doxycycline-inducible *NLRP3* group#3 variants were treated with doxycycline (1 μg/ml, 3h), LPS (40 ng/ml, 2h) and nigericin (15 μg/ml) in the presence of MCC950 (1 μM), CY-09 (50 μM), G5 (1 μM) and CRT006101 (0.5 μM) NLRP3 inhibitors or DMSO vehicle (added 20 min before the last treatment), and cell death was monitored by PI incorporation over time quantified by time-lapse high content microscopy for 2h. Means of area under the curve of duplicates and 1SD are represented. Two-way ANOVA multiple comparisons of each inhibitor with the corresponding treatment with vehicle only. *, p <0.05; **, p <0.01; ***, p <0.001; ****, p <0.0001. One experiment done in duplicates representative of 2 independent experiments is shown. Statistical analysis including all independent experiments are represented in Figure 7.

**Supplemental Figure 10.**
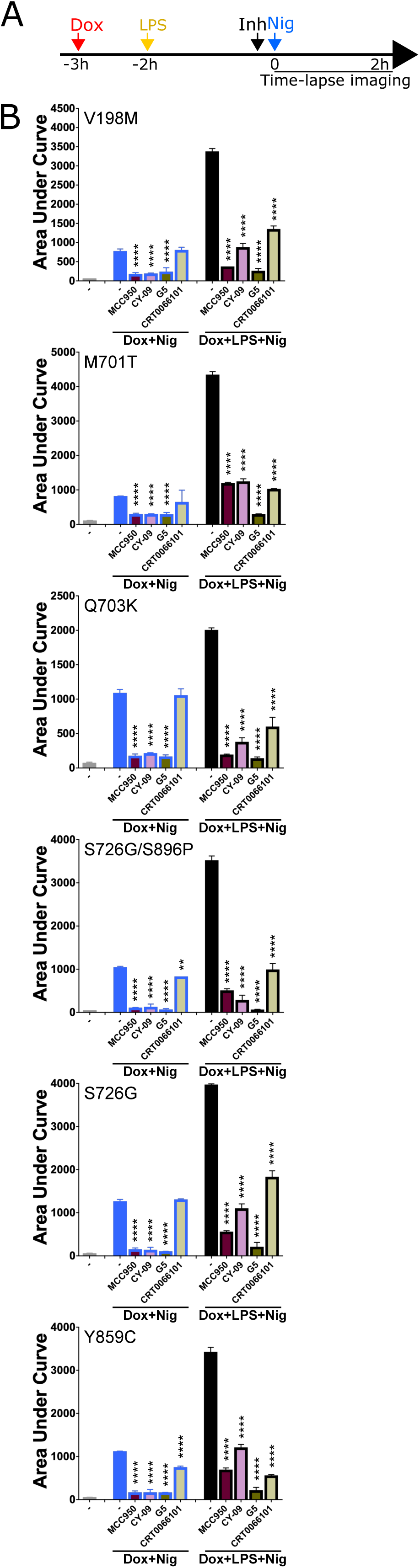
Sensitivity of group#4 *NLRP3* variants to NLRP3 inhibitors. A-B. *NLRP3*-deficient U937 cells reconstituted with doxycycline-inducible *NLRP3* group#4 variants were treated with doxycycline (1 μg/ml, 3h), LPS (40 ng/ml, 2h) and/or nigericin (15 μg/ml) in the presence of MCC950 (1 μM), CY-09 (50 μM), G5 (1 μM) and CRT006101 (0.5 μM) NLRP3 inhibitors or DMSO vehicle (added 20 min before nigericin), and cell death was monitored by PI incorporation over time quantified by time-lapse high content microscopy for 2h. Means of area under the curve of duplicates and 1SD are represented. Two-way ANOVA multiple comparisons of each inhibitor with the corresponding treatment with vehicle only. *, p <0.05; **, p <0.01; ***, p <0.001; ****, p <0.0001. One experiment done in duplicates representative of 2 independent experiments is shown. Statistical analysis including all independent experiments are represented in Figure 7.

**Supplemental Figure 11.**
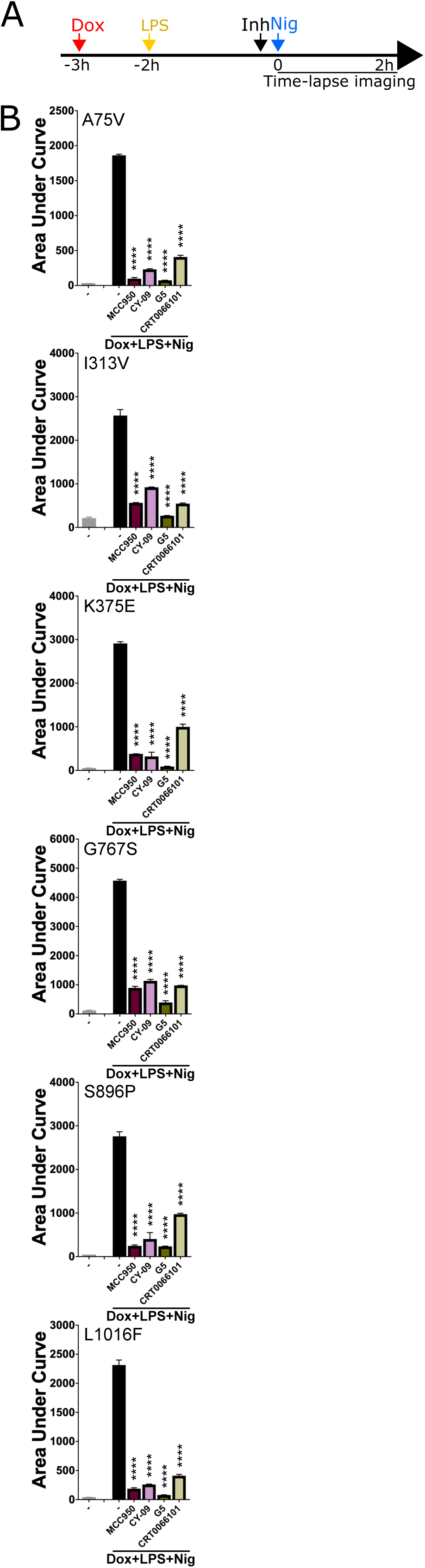
Sensitivity of group#5 *NLRP3* variants to NLRP3 inhibitors. A-B. *NLRP3*-deficient U937 cells reconstituted with doxycycline-inducible *NLRP3* group#5 variants were treated with doxycycline (1 μg/ml, 3h), LPS (40 ng/ml, 2h) and nigericin (15 μg/ml) in the presence of MCC950 (1 μM), CY-09 (50 μM), G5 (1 μM) and CRT006101 (0.5 μM) NLRP3 inhibitors or DMSO vehicle (added 20 min before nigericin), and cell death was monitored by PI incorporation over time quantified by time-lapse high content microscopy for 2h. Means of area under the curve of duplicates and 1SD are represented. Two-way ANOVA multiple comparisons of each inhibitor with the corresponding treatment with vehicle only. *, p <0.05; **, p <0.01; ***, p <0.001; ****, p <0.0001. One experiment done in duplicates representative of 2 independent experiments is shown. Statistical analysis including all independent experiments are represented in Figure 7.

**Supplemental Figure 12.**
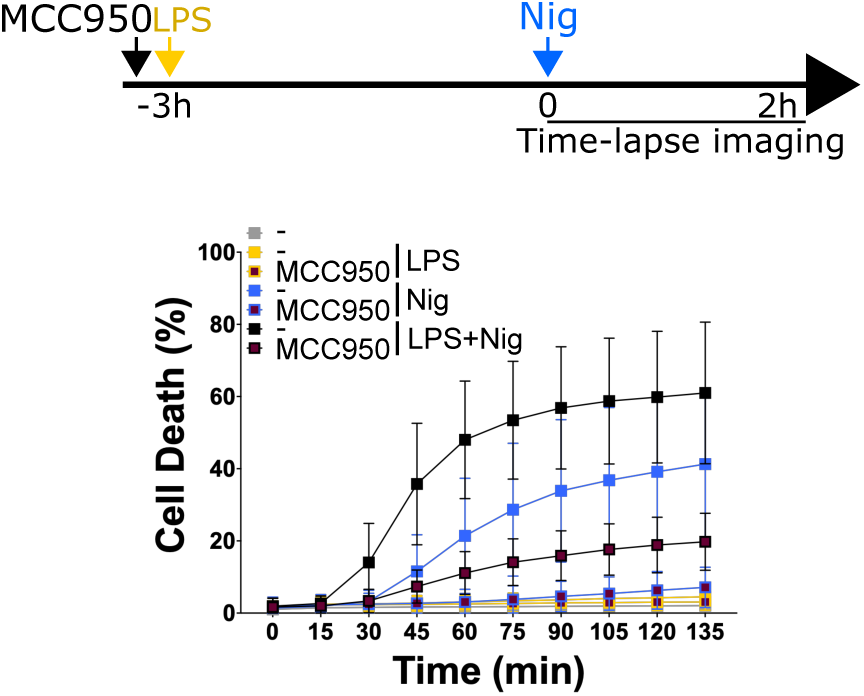
Nigericin activation, but not LPS priming, causes pyroptosis in primary monocytes from healthy donor. Monocytes of healthy donors (n=10) were treated with LPS (40 ng/ml, 3h) and nigericin (5 μg/ml) in the presence of MCC950 (1 μM, added 15 min before LPS) and cell death was monitored by PI incorporation over time quantified by time-lapse high content microscopy for 2h. Means and 1 SD are shown. 10 independent experiments with 1 donor each, done in 2-6 technical replicates are shown.

**Supplemental Figure 13.**
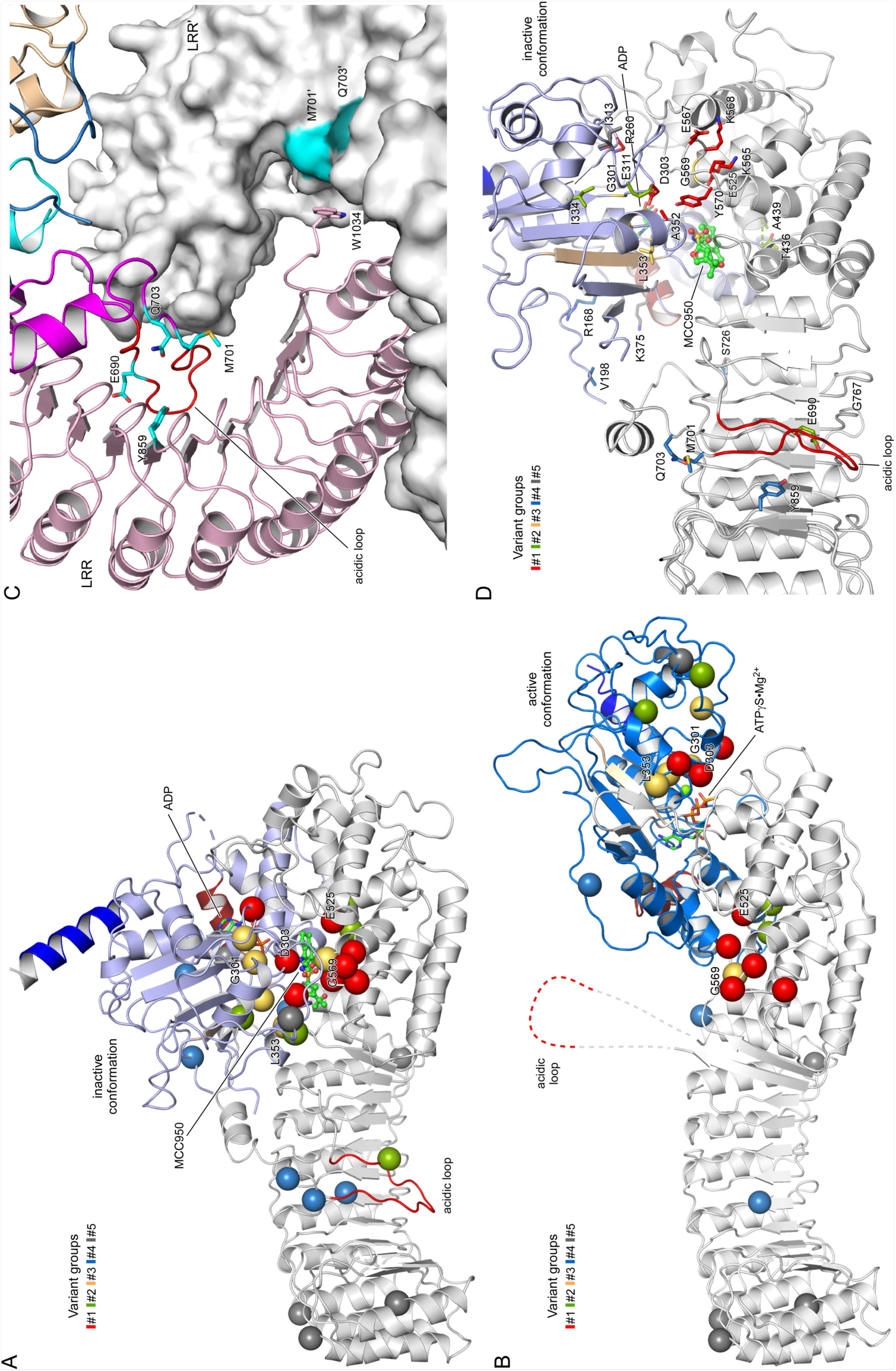
Details of the disease-causing mutant sites in NLRP3 and MCC950 inhibition. A, Localization of gain-of-function mutations in the MCC950-inhibited state of human NLRP3 (7PZC), color coded according to variant groups. The five mutant positions that are not responsive to MCC950 treatment, G301D, E303H, L353P, E525K, and G569R, are labelled. B, Localization of gain-of-function mutations in the adenosine triphosphate-bound active state of human NLRP3 (8EJ4). C, The three gain-of-function mutations in the acidic loop of NLRP3, E690, M701, and Q703, are in the dimer interface of the interlaced LRRs, while Y859 in the concave surface of the LRR interacts with the acidic loop. D, MCC950 is in the center of the NACHT and LRR domains in proximity to many disease-causing mutations.

## Notes

### Competing Interest Statement

The authors have declared no competing interest.

## References

1. Hoffman, H. M., Mueller, J. L., Broide, D. H., Wanderer, A. A. & Kolodner, R. D. Mutation of a new gene encoding a putative pyrin-like protein causes familial cold autoinflammatory syndrome and Muckle-Wells syndrome. Nat Genet 29, 301–5 (2001).

2. Booshehri, L. M. & Hoffman, H. M. CAPS and NLRP3. J. Clin. Immunol. 39, 277–286 (2019).

3. Louvrier, C. et al. NLRP3-associated autoinflammatory diseases: Phenotypic and molecular characteristics of germline versus somatic mutations. J. Allergy Clin. Immunol. 145, 1254–1261 (2020).

4. Van Gijn, M. E. et al. New workflow for classification of genetic variants’ pathogenicity applied to hereditary recurrent fevers by the International Study Group for Systemic Autoinflammatory Diseases (INSAID). J. Med. Genet. 55, 530– 537 (2018).

5. Bauernfeind, F. G. et al. Cutting edge: NF-kappaB activating pattern recognition and cytokine receptors license NLRP3 inflammasome activation by regulating NLRP3 expression. J Immunol 183, 787–91 (2009).

6. Juliana, C. et al. Non-transcriptional priming and deubiquitination regulate NLRP3 inflammasome activation. J Biol Chem 287, 36617–22 (2012).

7. McKee, C. M. & Coll, R. C. NLRP3 inflammasome priming: A riddle wrapped in a mystery inside an enigma. J. Leukoc. Biol. 108, 937–952 (2020).

8. Niu, T. et al. NLRP3 phosphorylation in its LRR domain critically regulates inflammasome assembly. Nat. Commun. 12, 5862 (2021).

9. Zhang, Z. et al. Distinct changes in endosomal composition promote NLRP3 inflammasome activation. Nat. Immunol. 24, 30–41 (2023).

10. Cuisset, L. et al. Mutations in the autoinflammatory cryopyrin-associated periodic syndrome gene: epidemiological study and lessons from eight years of genetic analysis in France. Ann Rheum Dis 70, 495–9 (2011).

11. Lagrange, B. et al. Human caspase-4 detects tetra-acylated LPS and cytosolic Francisella and functions differently from murine caspase-11. Nat. Commun. 9, 242 (2018).

12. Coll, R. C. et al. A small-molecule inhibitor of the NLRP3 inflammasome for the treatment of inflammatory diseases. Nat. Med. 21, 248–255 (2015).

13. Jiang, H. et al. Identification of a selective and direct NLRP3 inhibitor to treat inflammatory disorders. J. Exp. Med. 214, 3219–3238 (2017).

14. Hochheiser, I. V. et al. Structure of the NLRP3 decamer bound to the cytokine release inhibitor CRID3. Nature (2022) doi:10.1038/s41586-022-04467-w.

15. Brinkschulte, R. et al. ATP-binding and hydrolysis of human NLRP3. *Commun*. Biol. 5, 1176 (2022).

16. Tapia-Abellán, A. et al. MCC950 closes the active conformation of NLRP3 to an inactive state. Nat. Chem. Biol. 15, 560–564 (2019).

17. Dekker, C. et al. Crystal Structure of NLRP3 NACHT Domain With an Inhibitor Defines Mechanism of Inflammasome Inhibition. J. Mol. Biol. 433, 167309 (2021).

18. Py, B. F., Kim, M. S., Vakifahmetoglu-Norberg, H. & Yuan, J. Deubiquitination of NLRP3 by BRCC3 critically regulates inflammasome activity. Mol Cell 49, 331–8 (2013).

19. Zhang, Z. et al. Protein kinase D at the Golgi controls NLRP3 inflammasome activation. J Exp Med 214, 2671–2693 (2017).

20. Vande Walle, L., et al. MCC950/CRID3 potently targets the NACHT domain of wild-type NLRP3 but not disease-associated mutants for inflammasome inhibition. PLoS Biol. 17, e3000354 (2019).

21. Gaidt, M. M. et al. Human Monocytes Engage an Alternative Inflammasome Pathway. Immunity 44, 833–846 (2016).

22. Gritsenko, A. et al. Priming Is Dispensable for NLRP3 Inflammasome Activation in Human Monocytes In Vitro. Front. Immunol. 11, 565924 (2020).

23. Rieber, N. et al. A functional inflammasome activation assay differentiates patients with pathogenic NLRP3 mutations and symptomatic patients with low penetrance variants. Clin. Immunol. Orlando Fla 157, 56–64 (2015).

24. Theodoropoulou, K. et al. Increased Prevalence of NLRP3 Q703K Variant Among Patients With Autoinflammatory Diseases: An International Multicentric Study. Front. Immunol. 11, 877 (2020).

25. Xiao, L., Magupalli, V. G. & Wu, H. Cryo-EM structures of the active NLRP3 inflammasome disc. Nature 613, 595–600 (2023).

26. Andreeva, L. et al. Full-length NLRP3 forms oligomeric cages to mediate NLRP3 sensing and activation. bioRxiv 2021.09.12.459968 (2021) doi:10.1101/2021.09.12.459968.

27. Ohto, U. et al. Structural basis for the oligomerization-mediated regulation of NLRP3 inflammasome activation. Proc. Natl. Acad. Sci. U. S. A. 119, e2121353119 (2022).

28. Weber, A. N. R., et al. Effective ex vivo inhibition of cryopyrin-associated periodic syndrome (CAPS)-associated mutant NLRP3 inflammasome by MCC950/CRID3. Rheumatol. Oxf. Engl. 61, e299–e313 (2022).

29. Han, S. et al. Lipopolysaccharide Primes the NALP3 Inflammasome by Inhibiting Its Ubiquitination and Degradation Mediated by the SCFFBXL2 E3 Ligase. J Biol Chem 290, 18124–33 (2015).

30. Tang, J. et al. Sequential ubiquitination of NLRP3 by RNF125 and Cbl-b limits inflammasome activation and endotoxemia. J. Exp. Med. 217, (2020).

31. Wang, D. et al. YAP promotes the activation of NLRP3 inflammasome via blocking K27-linked polyubiquitination of NLRP3. Nat. Commun. 12, 2674 (2021).

32. Milhavet, F. et al. The infevers autoinflammatory mutation online registry: update with new genes and functions. Hum. Mutat. 29, 803–808 (2008).

33. Meerbrey, K. L. et al. The pINDUCER lentiviral toolkit for inducible RNA interference in vitro and in vivo. Proc. Natl. Acad. Sci. U. S. A. 108, 3665–3670 (2011).

